# Glycerol-Driven Energy and Proteostasis Underpin Antibiotic Tolerance in *Escherichia coli*

**DOI:** 10.64898/2026.05.19.725944

**Authors:** Han G. Ngo, Sayed Golam Mohiuddin, Mehmet A. Orman

## Abstract

Bacterial persisters are frequently described as metabolically dormant, yet the endogenous metabolic programs that sustain survival during prolonged nutrient limitation remain poorly understood. Here, using stationary-phase *Escherichia coli* as a model of antibiotic tolerance, we combine proteomics, genetics, metabolic phenotyping, and single-cell imaging to define a metabolic framework underlying persistence. Perturbation of tricarboxylic acid cycle function broadly reprogrammed stationary-phase physiology, suppressing lipid and glycerol metabolism, altering energy homeostasis and proteostasis, and reducing antibiotic tolerance. Systems-level analyses identified phospholipid-derived glycerol catabolism as a central metabolic node linking endogenous carbon recycling to persistence. Genetic disruption of glycerol utilization impaired proton motive force homeostasis, reduced formation of large polar protein aggregates, altered division-associated remodeling, and sensitized cells to antibiotic-induced lysis. Functional metabolic assays further revealed that persisters retain a selective capacity to utilize glycerol for rapid proton motive force restoration without growth resumption. Together, our findings support a model in which stationary-phase persisters are not metabolically inert but sustained through endogenous metabolic rewiring that coordinates energy maintenance, proteostasis, and antibiotic tolerance.

## Introduction

Although bacterial persisters have been discovered and studied for over eight decades ^1,2^, their underlying mechanisms are still elusive. This is largely due to the abundance and complexity of bacterial survival strategies, which involve numerous cellular processes that vary depending on environmental factors, growth and treatment conditions, antibiotic types, and bacterial species ^3–7^. Persisters have been identified in a wide range of bacteria, including *Escherichia coli*, *Mycobacterium tuberculosis*, and *Staphylococcus aureus*, as well as in fungi and even cancer cells ^3–11^. These cells are capable of surviving prolonged and intensive antibiotic or chemotherapeutic treatments without acquiring genetic mutations that induce resistance ^12^. Persisters are thought to be one of the reasons why antibiotic treatment failure, relapse of chronic infections, and the emergence of antibiotic resistance are happening ^13–16^. Despite significant progress, a comprehensive understanding of the diverse and context-dependent mechanisms of persistence is still needed to guide the development of effective anti-persister therapies.

Persister cells are frequently characterized as metabolically dormant. While this description captures an important aspect of their physiology (i.e., reduced anabolism and proliferation relative to exponentially growing cells), they should retain a basal level of metabolic activity and energy production ^17–20^. This basal metabolism may support key cellular functions such as resuming growth upon antibiotic removal ^21^, initiating stress responses like the SOS pathway ^22^, repairing DNA damage ^23^, sustaining energy through possible futile cycles ^7^, and utilizing specific carbon sources that potentiate aminoglycoside killing ^24^, all of which indicate a degree of physiological robustness in persisters. Indeed, more recent studies reveal that persister cells are highly metabolically heterogeneous ^25^ and can dynamically reprogram gene expression and regulatory networks to survive antibiotic stress ^26^.

Our previous work demonstrated that cyclic adenosine monophosphate (cAMP), together with its receptor protein CRP, functions as a key metabolic regulator during the stationary phase, shifting cellular metabolism from anabolic processes toward oxidative phosphorylation ^27^. This metabolic transition supports a basal level of energy production, which may be required for the survival of persister cells formed under these conditions ^27^. Consistent with this idea, genetic disruption of these regulatory and metabolic pathways significantly reduced persistence, particularly against ampicillin and ofloxacin ^27^. For instance, in our *E. coli* Keio knockout screen, we observed the most substantial reductions in persister levels among strains lacking key genes, always involved in the tricarboxylic acid (TCA) cycle, electron transport chain (ETC), and ATP synthase, while a few gene deletions in TCA and ETC components (such as *icd*, *nuoL*) led to modest increases in persistence ^27^. While these findings were further validated in independently constructed *E. coli* MG1655 deletion strains ^27^, the factors mediating these pathways in the stationary phase, particularly carbon sources or intracellular recycling, remain to be determined.

Bacterial cells in the stationary phase, particularly in nonnutritive environments, initiate intracellular self-digestion by degrading their own components, such as proteins and lipids, which serves as a survival strategy under nutrient-limiting conditions ^28–31^. Our previous work showed that this intracellular degradation correlates with persistence, likely by supplying essential metabolites and energy precursors ^32^, as we found that inhibiting stationary-phase metabolism simultaneously reduces both intracellular degradation and persistence ^32^. In eukaryotes, autophagy is a well-established mechanism for maintaining viability under nutrient deprivation and has been directly linked to drug tolerance. By contrast, analogous processes of self-digestion and intracellular recycling in bacteria remain far less well defined. In particular, the specific metabolic pathways and molecular players, including preferred carbon sources, driving this self-sustaining metabolic state remain largely uncharacterized. Resolving this knowledge gap is essential to understanding the metabolic underpinnings of antibiotic tolerance. In this study, we began with proteomic analysis of the succinate dehydrogenase knockout (Δ*sdhA*), a strain exhibiting markedly reduced stationary-phase persisters ^27^, to pinpoint metabolic pathways most disrupted in the stationary phase when persistence is lost. We systematically investigated the role of these metabolic components in shaping persister physiology and found that stationary-phase *Escherichia coli* can recycle lipids to fuel glycerol metabolism, maintain energy, and survive antibiotics.

## Results

### SdhA deletion rewires stress, redox, and metabolic pathways in stationary-phase E. coli

To investigate the physiological consequences of energy metabolism disruption in persister formation, we performed untargeted proteomic analysis on the Δ*sdhA* mutant during the late stationary phase. To reach this phase, overnight cultures were diluted into Luria-Bertani (LB) medium and grown for 24 hours, following the same experimental strategy used in our previous studies ^27,32^. This stage is critical for persistence, as cells typically transition into a low-metabolic-activity, stress-adapted state that promotes survival under antibiotic exposure. Δ*sdhA* was chosen because it represents a central node in both the TCA cycle and ETC, and consistently yields fewer persisters when transferred from stationary phase to fresh medium under antibiotic treatment ^27,33^. We first assessed global protein abundance in the Δ*sdhA* knockout and compared it to the wild-type (WT) strain using a volcano plot (**Figure 1a**, and **Supplementary Proteomics Data**), which illustrates fold-changes in protein expression. Statistical significance was assessed using a parametric t-test, with an adjusted p-value cutoff of 0.05 ^34^. The functions of significantly upregulated and downregulated proteins are listed in **Supplementary Table S1**. Proteins showing at least a two-fold increase or decrease in abundance were then analyzed using STRING-based network and pathway enrichment tools (**Figures 1b-e**), incorporating curated databases such as Gene Ontology, KEGG pathways, and UniProt functional terms ^35^. Enrichment strength was calculated as the log_10_ ratio observed to expected proteins per pathway, and significance was determined using a false discovery rate-corrected p-value threshold ^35^.

**Figure 1.**
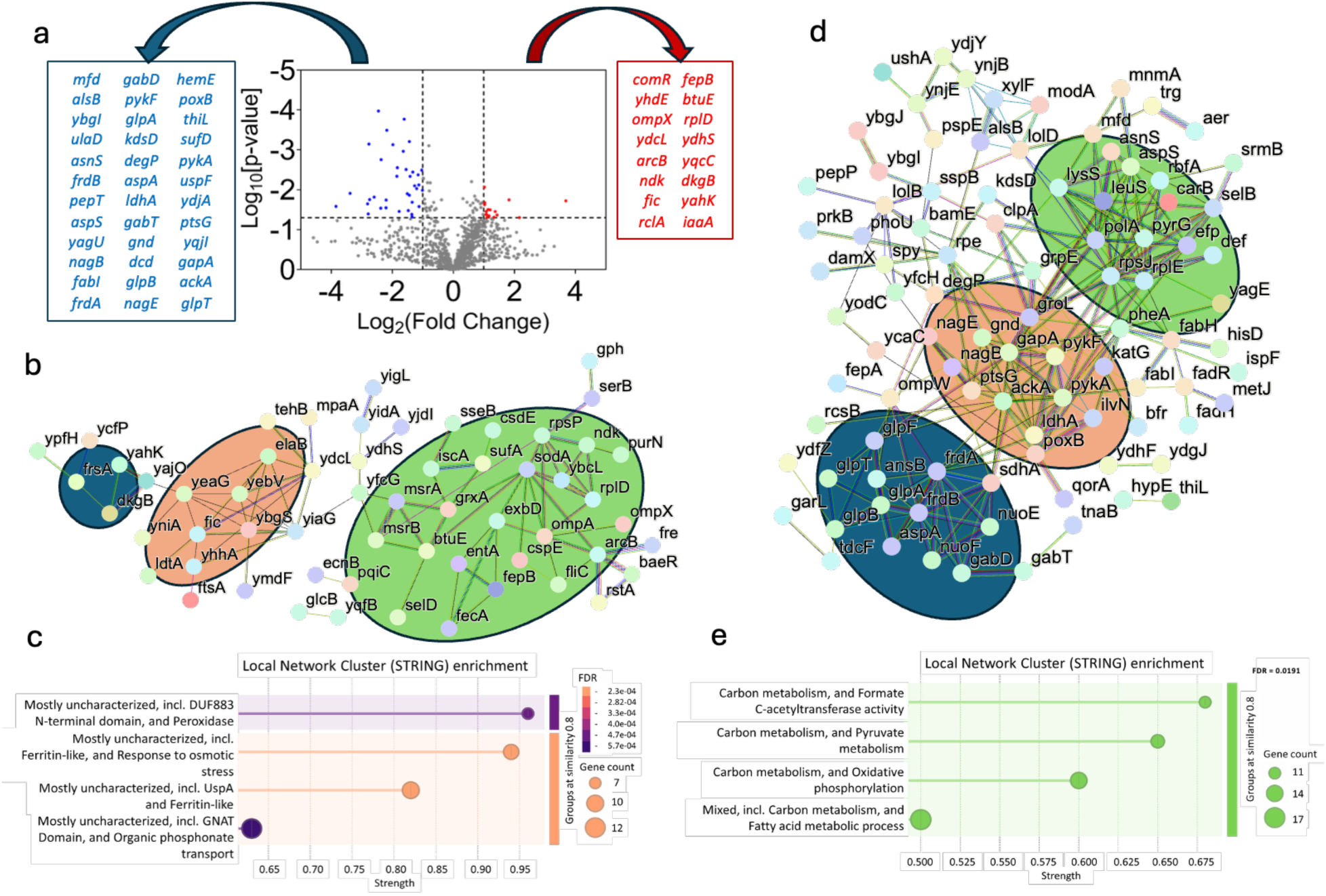
Proteomics analysis of the *sdhA* knockout strain during late stationary phase. (a) Volcano plot showing significantly upregulated (red) and downregulated (blue) proteins. The fold-change thresholds were set at ≤ –2 and ≥ 2, with a p-value cutoff of 0.05. (b) Pathway enrichment analysis of upregulated protein networks using STRING ^35^. Observed clusters are highlighted with colored circles: blue, detoxification and redox balance; orange, stringent response and stationary-phase adaptation; green, oxidative stress response, iron homeostasis, and global stress signaling. (c) Local network cluster enrichment (STRING) for the upregulated protein sets. (d) Pathway enrichment analysis of downregulated protein networks using STRING ^35^. Observed clusters are highlighted with colored circles: blue, oxidative phosphorylation, glycerol metabolism, and electron transport; orange, central carbon metabolism including glycolysis and pyruvate processing; green, translation and protein synthesis. (e) Local network cluster enrichment (STRING) for the downregulated protein sets. The STRING network illustrates protein-protein interactions in “evidence” mode, with edge colors indicating the type of supporting evidence, including curated databases, experimental data, gene neighborhood, gene fusions, co-occurrence, co-expression, protein homology, and text mining. Each local network cluster includes a false discovery rate (FDR), representing p-values adjusted for multiple testing using the Benjamini–Hochberg procedure. “Strength” denotes the enrichment effect, calculated as log_10_(observed/expected). The number of biological replicates for all panels, N = 3.

The network of upregulated proteins in Δ*sdhA* (**Figure 1b, c**) revealed three major functional clusters associated with (i) detoxification and redox balance, (ii) stringent response and stationary-phase adaptation, and (iii) oxidative stress response, iron homeostasis, and global stress signaling. The first group included oxidoreductases such as YahK and DkgB, which are involved in detoxifying reactive aldehydes and methylglyoxal, as well as YajO, a thiamine biosynthesis enzyme (the blue cluster, **Figure 1b**). These proteins likely mitigate damage from metabolic byproducts and contribute to maintaining redox homeostasis ^36^. The second cluster comprised proteins associated with the stringent response and stress adaptation, including YeaG, Fic, ElaB, YiaG, and others (the orange cluster, **Figure 1b**). These proteins are known to be induced under nutrient limitation and stress and are often regulated by the alarmone ppGpp ^37^. For example, Fic is a stationary-phase regulatory protein involved in energy sensing, and ElaB contributes to survival under oxidative and membrane stress ^38,39^. The third and most extensive group of upregulated proteins consisted of enzymes and regulators involved in oxidative stress defense (MsrA, MsrB, BtuE, GrxA, SodA, Fre), iron-sulfur cluster assembly (IscA, SufA), and iron transport (EntA, FepB, FecA, ExbD) (the green cluster, **Figure 1b**). Expression of these proteins indicates a broad cellular response to oxidative burden and iron limitation, conditions typical of the stationary phase, and interestingly, they are significantly elevated in Δ*sdhA*. Although one might expect Δ*sdhA* to reduce oxidative stress, the upregulation of these defense mechanisms may play a protective role, potentially contributing to the maintenance of proteostasis. Also, several components of the ribosome (RpsB, RplD), amino acid metabolism (SerB), and nucleotide balance (Ndk) were upregulated (**Figure 1b, c**), possibly reflecting the need to maintain core metabolic functions despite energy stress. Additionally, regulators such as ArcB, BaeR, and RstA, which mediate anaerobic and envelope stress responses ^40–42^, were more abundant, suggesting transcriptional reprogramming in response to metabolic disruption.

In contrast, the set of downregulated proteins (**Figure 1d, e**) pointed to a repression of core cellular functions in Δ*sdhA*. We identified three major clusters of functionally related proteins associated with (i) translation and protein synthesis, (ii) central carbon metabolism including glycolysis and pyruvate processing, and (iii) oxidative phosphorylation, lipid degradation and glycerol metabolism. Reduced abundance of ribosomal proteins (RpsJ, RplE, MnmA, AsnS, AspS) and translation factors (Efp, SelB, Def) (the green cluster, **Figure 1d**) suggests a general slowdown in protein synthesis, consistent with energy limitation in stationary-phase cells lacking a functional TCA cycle. Key glycolytic and associated enzymes, including GapA, PykF, Gnd, LdhA, PoxB, and PtsG, were also significantly downregulated (the orange cluster, **Figure 1d**). These enzymes are responsible for carbon flux through glycolysis and pyruvate metabolism and are tightly linked to energy generation and biosynthetic precursor supply ^37^. Thiamine pyrophosphate (TPP)-dependent enzymes and TPP-binding proteins were especially affected, indicating that decarboxylation steps and cofactor metabolism may be impaired in Δ*sdhA*. Further repression was observed in proteins involved in oxidative phosphorylation and anaerobic respiration (the blue cluster, **Figure 1d**). In addition to the expected loss of SdhA, components of fumarate and NADH dehydrogenase complexes (FrdA, FrdB, NuoE, NuoF) were also decreased. These changes are likely to restrict the cell’s capacity to generate ATP and intermediates necessary for biosynthesis. Multiple proteins involved in glycerol utilization and alternative carbon source metabolism (GlpA, GlpB, GlpF, and YdfZ) were downregulated, suggesting that the mutant is unable to efficiently reroute carbon flux under nutrient-limited conditions. STRING enrichment analysis revealed additional downregulation of pathways such as fatty acid metabolism and formate C-acetyltransferase activity (**Figure 1d,e**), supporting the notion that the energy and biosynthetic flexibility required for stationary-phase survival is compromised in the Δ*sdhA* background. Overall, these results demonstrate that *sdhA* deletion induces broad metabolic and stress-response reprogramming that reshapes stationary-phase physiology, with these alterations further verified in the subsequent sections.

### TCA cycle knockouts do not exhibit altered oxidative damage in the stationary phase

Although our proteomic analysis focused on the *ΔsdhA* knockout, the low-persister phenotype and associated metabolic shifts observed in this strain reflect broader physiological trends among other TCA cycle mutants (e.g., *Δlpd*, *ΔgltA*, *ΔsucC*, *Δmdh*), as most of these knockouts generate significantly fewer persisters than WT upon transfer from the stationary phase to fresh medium (thus, these results are not repeated here) ^27,32^. To ensure that these phenotypes were not confounded by major growth abnormalities, we first examined the growth dynamics of the TCA cycle knockouts. Growth curves based on optical density (OD_600_) measurements in LB medium showed that all tested knockouts exhibited growth similar to the WT, except for Δ*lpd* (**Figure 2a**). Flow cytometry–based cell counts at the late stationary phase (24 hours), which provide a more accurate measure of cell abundance, further confirmed that final cell densities were generally comparable across all strains, again with the exception of Δ*lpd* (**Supplementary Figure S1**). While slow growth is thought to be associated with increased persistence, the Δ*lpd* strain challenges this assumption, as it exhibits reduced persistence ^27^, despite its growth defect. Since most mutants do not exhibit growth deficiencies, and all show reduced persistence ^27^, these findings suggest that the low-persister phenotype and downstream phenotypic differences are not attributable to impaired growth or reduced viability in stationary phase. Of note, all follow-up assays were performed using normalized cell numbers to ensure consistency across strains.

**Figure 2.**
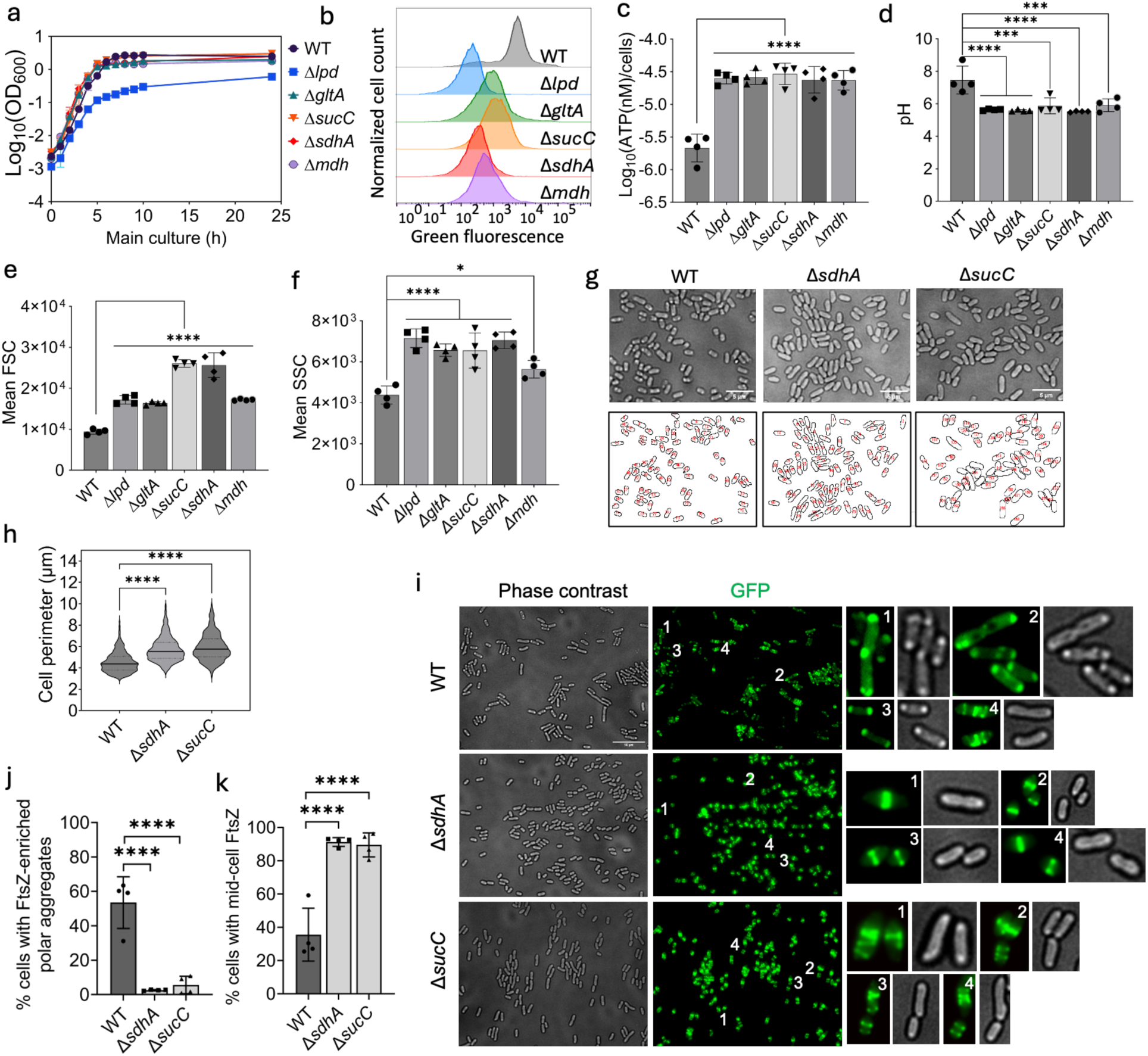
Physiological and morphological characterization of TCA cycle knockout strains in late stationary phase. (a) Growth curves of wild-type and TCA cycle knockout strains measured by OD_600_ (N = 3). (b) Redox activity of wild-type and mutant strains assessed using RedoxSensor Green (RSG) dye and analyzed by flow cytometry (N = 3). A representative plot is shown; all biological replicates produced similar results. (c) ATP levels measured using the BacTiter-Glo Microbial Cell Viability Assay (N = 4). (d) Intracellular pH levels measured using pGFPR01, which expresses the ratiometric fluorophore pHluorin (N = 4). (e, f) Forward scatter (FSC) and side scatter (SSC) measurements obtained by flow cytometry (N = 4). (g) Phase-contrast micrographs of wild-type and mutant strains. A representative image is shown; similar results were observed across four biological replicates (N = 4). Scale bar: 5 μm. Cell perimeters were quantified using ImageJ, with segmented micrographs shown below. (h) Quantification of cell perimeters from 1,000 cells. (i) Microscope imaging to visualize structural organization of Z-rings and large polar aggregates in *E. coli* WT, Δ*sdhA*, and Δ*sucC* strains. Representative phase-contrast and corresponding fluorescence images are shown, with selected cells (numbered) enlarged and displayed side by side to highlight diverse morphologies and structural features. (N = 4). All biological replicates produced similar results. Scale bar, 10 µm. (j) Quantification of the percentage of cells containing large polar aggregates (LPAs) in *E. coli* WT, Δ*sdhA*, and Δ*sucC* strains. Each dot represents the mean of a biological replicate, based on analysis of thousands of cells across multiple images (N = 4). (k) Quantification of the percentage of cells containing Z-rings in *E. coli* WT, Δ*sdhA*, and Δ*sucC* strains. Each dot represents the mean of a biological replicate, based on analysis of thousands of cells across multiple images (N = 4). All statistical analyses were performed using one-way ANOVA with Dunnett’s multiple comparisons test. Statistical significance is indicated as follows: *P <0.05; ***P < 0.001; ****P < 0.0001. Data represent the mean ± standard deviation.

Given that many of the proteomic changes in Δ*sdhA* were associated with oxidative stress and redox homeostasis, we next examined whether loss of TCA cycle enzymes results in altered oxidative damage. First, we assessed cell membrane integrity using propidium iodide (PI) staining and flow cytometry. No significant differences in membrane damage were observed between wild-type and knockout strains at the late stationary phase (**Supplementary Figure S2**). To more directly evaluate oxidative stress, we measured three well-established markers of oxidative damage: DNA oxidation, protein carbonylation, and lipid peroxidation (**Supplementary Figure S3**). DNA oxidation was quantified using an ELISA-based assay targeting 8-hydroxy-2’-deoxyguanosine and related oxidized guanine species, following enzymatic digestion of DNA extracted from stationary-phase cultures ^43^. Protein carbonylation was measured using a colorimetric assay based on 2,4-dinitrophenylhydrazine (DNPH), which reacts with carbonyl groups on oxidatively modified proteins to form detectable hydrazone derivatives ^44^. Lipid peroxidation was assessed by quantifying malondialdehyde (MDA) levels using the thiobarbituric acid reactive substances (TBARS) assay. MDA is a stable end-product of lipid peroxidation and serves as a reliable marker of oxidative damage to membrane lipids ^45^. No significant differences in oxidative DNA damage, protein carbonylation, or lipid peroxidation were detected in any of the TCA cycle knockouts compared to the wild type (**Supplementary Figure S3**). Thus, although TCA cycle disruption does not measurably affect oxidative damage, the redox response revealed by our proteomics data (**Figure 1**) likely reshapes proteostasis. Also, the reduced persistence of TCA cycle knockouts is unlikely to result from oxidative damage under these conditions, consistent with our previous findings ^32^.

### TCA cycle disruption reprograms stationary-phase energy homeostasis

Our proteomic analysis of the Δ*sdhA* mutant revealed pronounced changes in oxidative phosphorylation, proteostasis, and lipid degradation pathways, pointing to broad remodeling of stationary-phase physiology. Guided by these findings, we evaluated corresponding phenotypic parameters, including metabolic activity, intracellular ATP levels, cytoplasmic pH, cell size and complexity, and protein aggregation. We first used Redox Sensor Green (RSG) dye, which fluoresces in response to cellular reductase activity and serves as a general indicator of bacterial metabolic activity (**Figure 2b**). Flow cytometry analysis of late stationary-phase cells revealed that all TCA cycle knockouts exhibited lower RSG fluorescence than WT (**Figure 2b**), consistent with our proteomics data indicating reduced metabolic activity (**Figure 1**).

ATP levels were measured in the late stationary phase using a luciferase-based assay, and interestingly, all knockouts exhibited significantly higher intracellular ATP concentrations compared to WT (**Figure 2c**). The observed ATP elevation persists throughout the stationary phase (**Supplementary Figure S4**). Although intriguing, inverse metabolic relationships in genetically perturbed strains, such as those with TCA cycle deletions or ATP synthase mutants, have also been documented in previous studies ^46,47^. We infer that the increase in ATP levels arises from reduced utilization rather than enhanced production in our study, as supported by proteomics demonstrating decreased TCA cycle and ETC activity alongside a broader metabolic downshift (**Figure 1**). If ATP synthase and respiratory activity are reduced in the knockout strains, we would expect impaired proton export and a potential shift toward fermentative metabolism (leading to the accumulation of organic acids), both of which contribute to cytoplasmic acidification.

Therefore, we next measured intracellular pH using the pH-sensitive GFP variant pHluorin, expressed from an inducible plasmid (**Figure 2d**). The pHluorin reporter has a dual excitation peak at 410 and 470 nm with an emission maximum at 530 nm ^48,49^. Changes in intracellular pH alter the excitation ratio, enabling ratiometric measurement ^48,49^. All TCA cycle knockouts displayed significant cytoplasmic acidification relative to WT in late stationary phase (**Figure 2d**), further supporting a global metabolic downshift in these strains and corroborating the findings of our proteomics analysis (**Figure 1**).

### TCA cycle disruption alters stationary-phase cellular architecture

Next, we employed flow cytometry to assess cell size and complexity using forward and side scatter signals, respectively (**Figure 2e, f**). During the stationary phase, self-digestion of proteins and membrane phospholipids typically drives cell shrinkage and loss of cytoplasmic content, reducing both parameters; and our previous work showed that inhibiting stationary-phase metabolism suppresses self-digestion ^32^. Consistent with this, TCA cycle knockouts generally showed increased forward scatter (FSC, **Figure 2e**), indicating larger cell size, and elevated side scatter (SSC, **Figure 2f**), reflecting greater intracellular complexity. These findings were validated by phase-contrast microscopy for selected knockout strains (**Figure 2g**). Visual inspection and quantitative cell perimeter (**Figure 2g**) analysis confirmed that knockout cells were significantly larger than WT cells in the stationary phase (**Figure 2h**). Notably, these effects were specific to the stationary phase, as no consistent trends were observed in FSC and SSC profiles during exponential growth (**Supplementary Figure S5**).

Microscopy also revealed differences in protein aggregation. WT cells frequently showed bright polar foci in phase contrast images, likely corresponding to large polar aggregates (LPAs), while these structures were less pronounced in the TCA cycle knockouts (**Figure 2g**). These aggregates are a hallmark of aging stationary-phase cells and have been associated with delayed regrowth and persister cell formation ^50–54^. To further investigate these LPAs quantitatively in stationary-phase cells, we employed a low-copy plasmid-based reporter system (pUA66-*ftsZ-gfp*), in which the *ftsZ* gene is fused to *gfp* and placed under the control of an IPTG-inducible T5 promoter (**Figure 2i**). FtsZ is an essential cytoskeletal protein that orchestrates bacterial cell division by forming the Z-ring at the division site ^55–58^. We selected FtsZ-GFP as a reporter for two key reasons: (i) under stress conditions, FtsZ has been shown to localize to protein aggregates, making it a suitable proxy for visualizing and quantifying these structures, and (ii) Z-ring formation is critical for the resumption of growth in dormant cells, directly linking FtsZ dynamics to persistence and antibiotic susceptibility ^50–53,55–60^.

To visualize Z-ring formation and protein aggregation, late stationary-phase cells from WT, Δ*sdhA*, and Δ*sucC* strains harboring the FtsZ reporter plasmid were transferred to phosphate-buffered saline (PBS) agarose pads and imaged using fluorescence microscopy (**Figure 2i)**. Based on the localization patterns of FtsZ-GFP, we observed highly heterogeneous structures, including well-formed single Z-rings, multiple Z-rings within the same cell, filamentous morphologies, FtsZ-enriched LPAs, mid-cell localization of FtsZ proteins, and in some cases, a complete absence of detectable FtsZ signal (**Figure 2i**). Nearly half of WT cells displayed clear FtsZ-enriched LPAs, whereas this phenotype was significantly reduced in both the Δ*sdhA* and Δ*sucC* mutants (**Figure 2j**). This analysis was performed on large sample sizes: 7,391 WT cells, 6,201 Δ*sdhA* cells, and 5,494 Δ*sucC* cells. In general, Z-ring–like structures were more frequently observed in the mutants compared to WT, although these were also heterogeneous, often appearing as multiple rings or as mid-cell FtsZ assemblies rather than polar foci (**Figure 2k**). Thus, despite the structural variability, a clear reduction in LPA formation was evident in the mutants relative to WT, indicating that loss of LPAs can be associated with reduced persistence in TCA cycle knockouts (**Figure 2i–k**).

To further investigate the relationship between persistence and LPAs, we monitored individual cells by time-lapse microscopy during antibiotic exposure. Late stationary-phase cultures were inoculated onto LB agarose pads containing ampicillin (200 µg/mL), sealed with coverslips, and imaged in an on-stage incubation chamber maintained at 37 °C. Cells were monitored for up to 5 hours, as longer time courses were limited by agarose pad deformation and loss of microscopy focus. Ampicillin was chosen for these experiments because it facilitates real-time visualization of cell death. Ampicillin-induced lysis was highly dynamic, with cells becoming spherical spheroplasts or protoplasts before rupturing, collapsing, and releasing their contents (**Movie 1**).

By contrast, untreated controls continued to elongate and divide normally (**Figure 3**). We observed that Δ*sdhA* cells, which exhibit markedly reduced LPAs, underwent nearly complete lysis within 3 hours of ampicillin treatment, with lytic events detectable as early as 40 minutes (**Figure 3b**). Similarly, Δ*sucC* cells were also rapidly lysed, with only a small fraction of intact cells remaining (**Figure 3c**). In contrast, WT cells exhibited delayed lysis, and a substantial fraction of survivors retained LPAs and protein aggregates up to 5 hours (**Figure 3a**). These single-cell observations provide direct mechanistic support for the elevated persister levels seen in WT relative to TCA cycle mutants, in line with our study ^27^. Although the connection between persistence and protein aggregation is well established in the literature, our finding that enhanced proteostasis, driven by metabolic perturbations in TCA cycle mutants, sensitizes cells to antibiotic-induced lysis represents, to our knowledge, a novel link between energy metabolism, protein quality control, and antibiotic tolerance.

**Figure 3.**
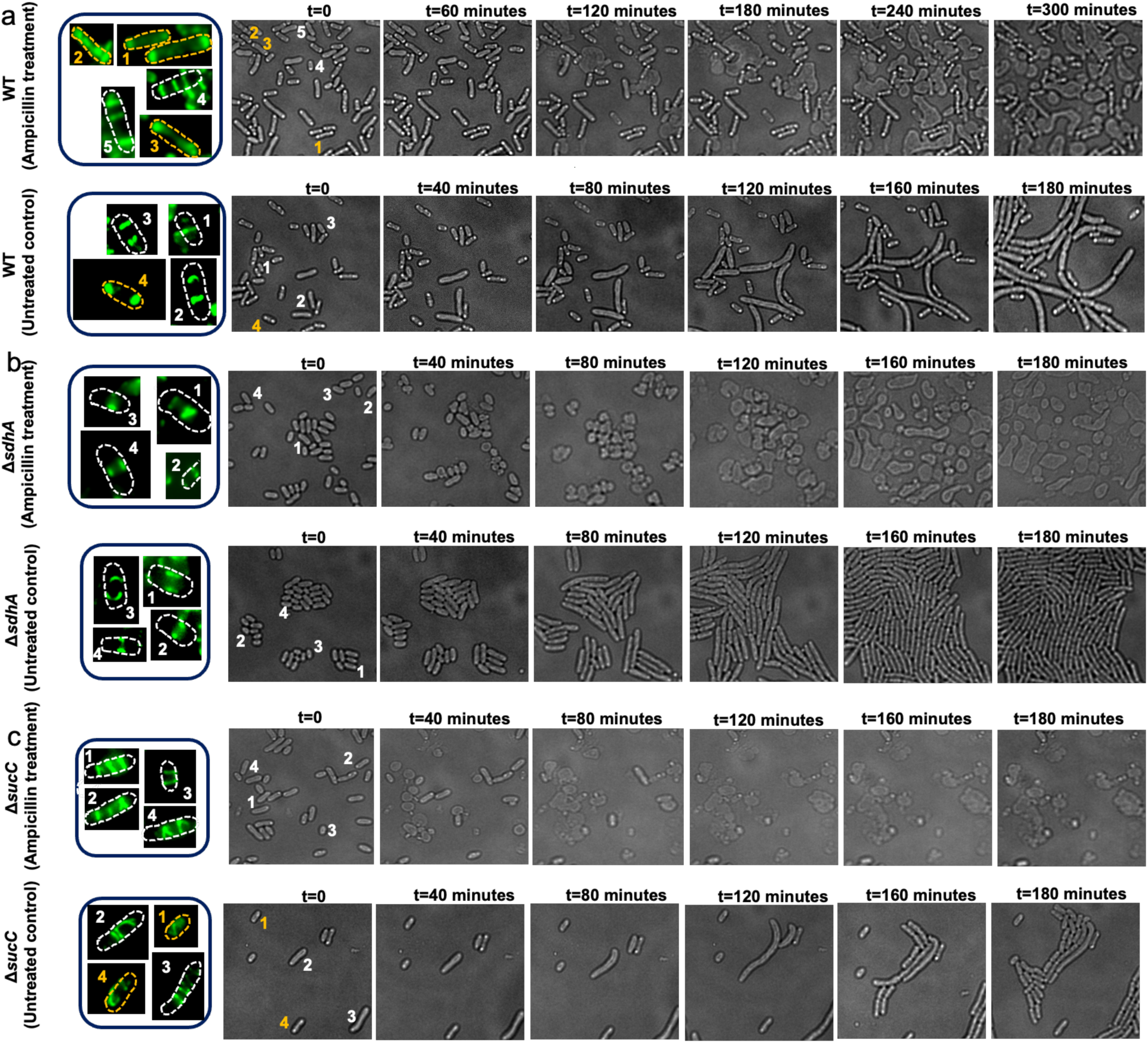
Time-lapse microscopy of (a) WT, (b) Δ*sdhA*, and (c) Δ*sucC* strains under untreated and ampicillin-treated conditions. Each panel contains two rows: ampicillin-treated cells (top) and untreated controls (bottom). In each row, the first column shows a fluorescent image of representative cells at t = 0 h (FtsZ–GFP), with numbered cells outlined by dotted lines to indicate orientation; orange marks non-growing or unlysed cells, and white marks growing or lysed cells. The second column shows the corresponding phase-contrast image at t = 0 h, and subsequent columns show phase-contrast images at the indicated time points. Fluorescent images were acquired only at t = 0 h to visualize Z-rings; repeated blue-laser excitation was avoided due to light sensitivity of the reporter and its potential to perturb growth and lysis dynamics. Images were collected at 37 °C in an on-stage incubator. Cell lysis or growth events can shift cell orientation and position during the time course. A representative image is shown; similar results were obtained across four biological replicates (N = 4).

Because overexpression of FtsZ–GFP can potentially perturb cell division, cellular size and architecture, and lysis dynamics, and because microscopy-based analyses sample a limited number of cells, we sought to independently validate antibiotic-induced lysis at the population level using an orthogonal approach. To this end, we employed a control reporter plasmid (pUA66-*gfp*) that uses the same vector backbone, promoter, and induction conditions as the FtsZ–GFP construct, but expresses GFP alone. We have extensively used this reporter, as well as closely related constructs, in our previous studies ^59,61–64^ to quantify cell integrity and antibiotic tolerance at the population level, where loss of GFP signal serves as a robust proxy for cell lysis, whereas GFP-retaining cells represent intact, non-lysed populations enriched for persisters and viable but nonculturable (VBNC) cells, as previously validated ^61,62^. Using this assay, stationary-phase cells expressing GFP were treated with ampicillin under the same conditions used for microscopy (**Figure 4**). Upon lysis, cells rapidly lost GFP fluorescence, whereas intact cells retained GFP (**Figure 4**). Consistent with our single-cell observations, wild-type cultures maintained a substantial fraction of GFP-positive intact cells, even after prolonged antibiotic exposure of up to 20 hours (**Figure 4a, Supplementary Figure S6**). In contrast, Δ*sucC* cultures showed a marked reduction in the fraction of intact GFP-positive cells, although a small subpopulation remained detectable (**Figure 4c, Supplementary Figure S6**). Expectedly, Δ*sdhA* cultures exhibited near-complete loss of GFP-positive cells, with intact cells becoming almost undetectable, closely mirroring the rapid and extensive lysis observed by time-lapse microscopy (**Figure 4b, Supplementary Figure S6**). Together, these population-level measurements corroborate our imaging data and further support the conclusion that disruption of TCA cycle–dependent metabolism profoundly sensitizes stationary-phase cells to antibiotic-induced lysis.

**Figure 4.**
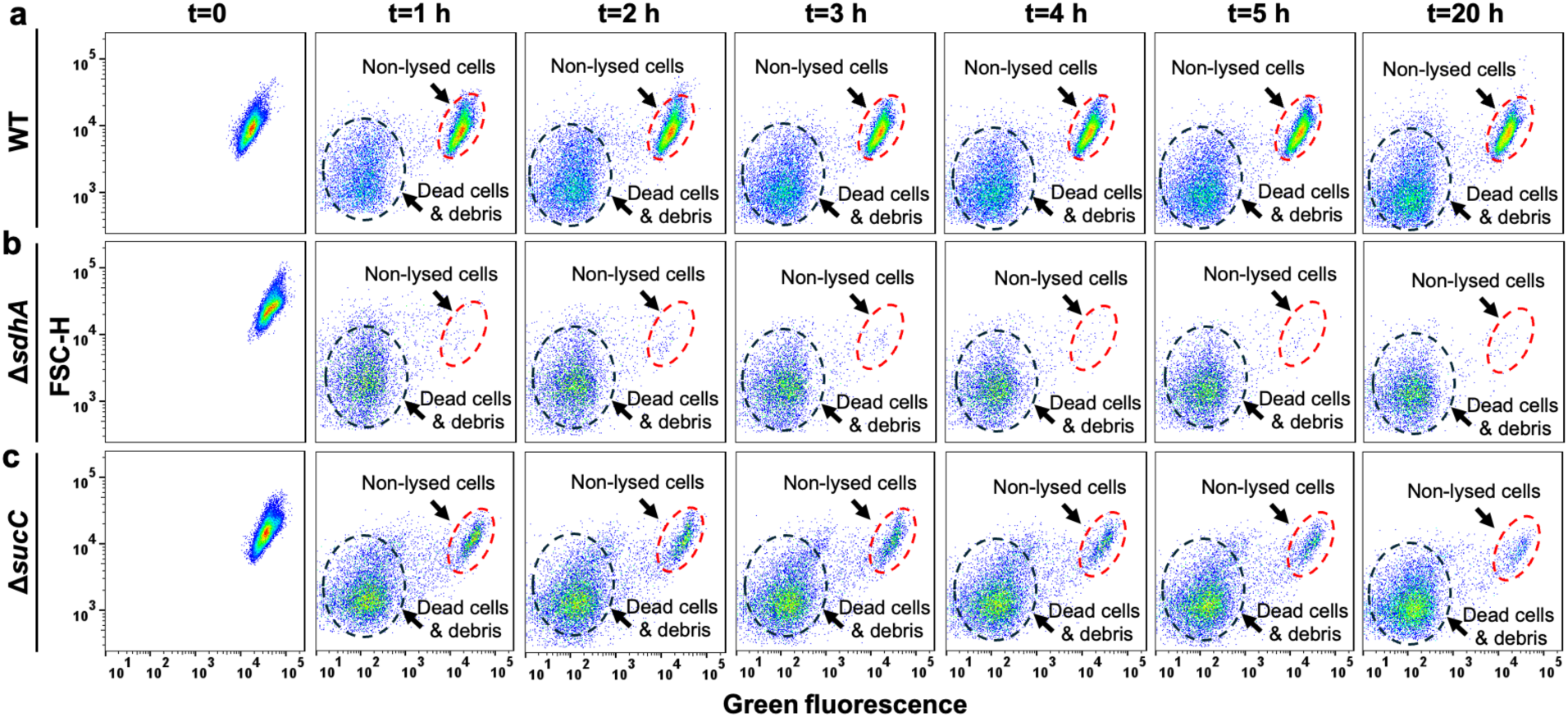
Flow cytometry analysis reveals reduced populations of intact cells in metabolic mutants. Cells harboring the GFP reporter plasmid pUA66-*gfp* were cultured in the presence of inducer (1 mM IPTG). At the late stationary phase, cultures were diluted 100-fold into fresh medium containing ampicillin and incubated under the same conditions used for microscopy assays. GFP fluorescence of individual cells was monitored by flow cytometry for (a) WT, (b) Δ*sdhA*, and (c) Δ*sucC*. Upon antibiotic-induced lysis, cells lose GFP signal, whereas intact cells retain high GFP fluorescence. Representative flow cytometry profiles are shown; all biological replicates produced similar results (N = 4).

Altogether, the results in **Figures 2, 3** and **4** demonstrate that disruption of the TCA cycle alters core aspects of stationary-phase physiology by suppressing metabolic activity, altering ATP levels, disrupting pH homeostasis, enlarging cell size, and preventing protein aggregation and mediating Z-rings. These changes underscore the TCA cycle’s role in cellular remodeling and proteostasis during stationary-phase adaptation. Notably, the differences in protein aggregation between WT and mutant strains are unlikely to result from oxidative damage (**Supplementary Figure S3**) but may instead reflect altered proteostasis associated with shifts in protein quality control, redox-balancing, and defense pathways in Δ*sdhA*, as highlighted by our proteomic analysis (**Figure 1**). Furthermore, the consistency between these metabolic and physiological observations (**Figure 2**) and our proteomic data (**Figure 1**) underlines the overall robustness and validity of our study.

### Keio knockout screening reveals key metabolic contributors to stationary-phase persistence

To investigate the molecular basis underlying reduced persistence in TCA cycle knockouts, particularly the Δ*sdhA* strain, we performed a targeted genetic screen using the Keio *E. coli* BW25113 knockout collection (**Figure 4a**). Although our proteomics analysis revealed major changes in stationary-phase physiology, the specific genes driving the low-persister phenotype remained unclear. To address this, we selected 50 key genes from the Δ*sdhA* proteomic dataset (**Supplementary Table S2**) and assessed the persistence levels of their corresponding knockout strains. We specifically focused on the key proteins identified in the upregulated and downregulated clusters highlighted in **Figure 1b** and **1d**, as these represent functional groups most likely to influence energy metabolism, stress adaptation, and proteostasis. For the experiments, equal numbers of stationary-phase cells were diluted in fresh media (∼100-fold) and treated for 20 hours with ampicillin (200 μg/mL) or ofloxacin (5 μg/mL) at levels well above the minimum inhibitory concentrations (MICs) and commonly used to study highly tolerant persister cells. Control strains included *lacY* and *lacI* knockouts, which are not expected to influence persistence^65^ but, like all Keio strains, carry a kanamycin resistance marker. Kanamycin was added to all cultures for selection and contamination control.

Our screen identified several knockouts with reduced persister levels, most corresponding to genes downregulated in the Δ*sdhA* proteome, including SelB, RbfA, Efp, CarB, GlpT, LysS, ClpA, AspA, GlpA, and PolA (**Figure 4a**). This supports the hypothesis that loss of these proteins contributes to the reduced persistence observed in Δ*sdhA*. While some knockouts showed elevated persistence (**Supplementary Table S2**), these were generally linked to genes upregulated in Δ*sdhA*. If deletion of these genes increases persistence, their upregulation in the mutant may act to counterbalance or limit persistence. Although these findings are intriguing and will be explored in future work, our current focus is on knockouts with reduced persistence, as they are most likely to reveal how metabolic disruption suppresses persistence and to provide direct functional links to the observed proteomic changes.

Many of these genes, whose knockouts reduced persistence, align with the physiological alterations observed in the TCA cycle knockouts. For instance, SelB, RbfA, and Efp are involved in translation and ribosome function, processes tightly linked to cell growth and stress adaptation (**Figure 1)**. The *clpA* gene encodes a key ATP-dependent protease involved in protein quality control and degradation, and its loss may contribute to the altered proteostasis and reduced protein degradation. Notably, *glpT* and *glpA* are central to glycerol metabolism, supplying intermediates to the TCA cycle as an alternative route ^37^. This pathway was found to be downregulated in the Δ*sdhA* proteome (**Figure 1**), indicating this route might be a critical factor of stationary-phase persistence.

### Glycerol catabolism sustains persistence in stationary-phase cells

We are particularly interested in glycerol metabolism because we hypothesize that stationary-phase cells in nutrient-depleted conditions catabolize phospholipids, generating glycerol that feeds into central metabolism to sustain PMF. While our initial analysis of proteomics data revealed broad metabolic changes (**Figure 1**), motivated by results from our Keio screen, we revisited our proteomic data with a specific focus on lipid-associated pathways (**Supplementary Proteomics Data**) and observed consistent downregulation of key proteins required for phospholipid degradation and glycerol processing in the Δ*sdhA* mutant. These include GlpF, GlpA, GlpB, GlpT, GlpK, TpiA, GlpQ, and the global regulator Crp (**Figure 5b**). These proteins mediate periplasmic degradation of glycerophosphodiesters to glycerol-3-phosphate (GlpQ); glycerol (GlpF) and periplasmic glycerol-3-phosphate transport (GlpT); phosphorylation and conversion into glycolytic intermediates (GlpK, TpiA); redox-linked processing (GlpA, GlpB); and transcriptional regulation under nutrient limitation (Crp) ^66^, which serves as the global regulator of this alternative glycerol metabolism ^37^. Furthermore, we re-examined two metabolomics datasets generated under comparable stationary-phase culture conditions in our previous studies and identified changes consistent with phospholipid-derived glycerol metabolism (**Supplementary Tables S3 and S4**) ^27,67^. Although those earlier analyses primarily focused on the energy metabolism of persisters rather than lipid pathways, the data clearly showed that major intracellular *E. coli* phospholipid components (which are abundant constituents of the cytoplasmic membrane) and glycerol-3-phosphate decreased at late stationary phase relative to early stationary phase ^27^, indicating active phospholipid utilization (**Supplementary Table S3**). In contrast, in cells treated with a PMF inhibitor, which experience collapsed PMF and reduced TCA and ETC activity similar to our metabolic mutants, the same lipids and glycerol-3-phosphate accumulated markedly ^67^, suggesting their limited utilization (**Supplementary Table S4**). While all of these data clearly point to the importance of phospholipid degradation (**Figure 5b**) and disruption of some of the key enzymes within them (e.g., GlpABC and GlpD, see **Figure 8a**) has also been shown by others to reduce persistence ^68^, whether glycerol-derived metabolites are directly used by stationary-phase persisters to sustain PMF and confer antibiotic tolerance remains unresolved.

**Figure 5.**
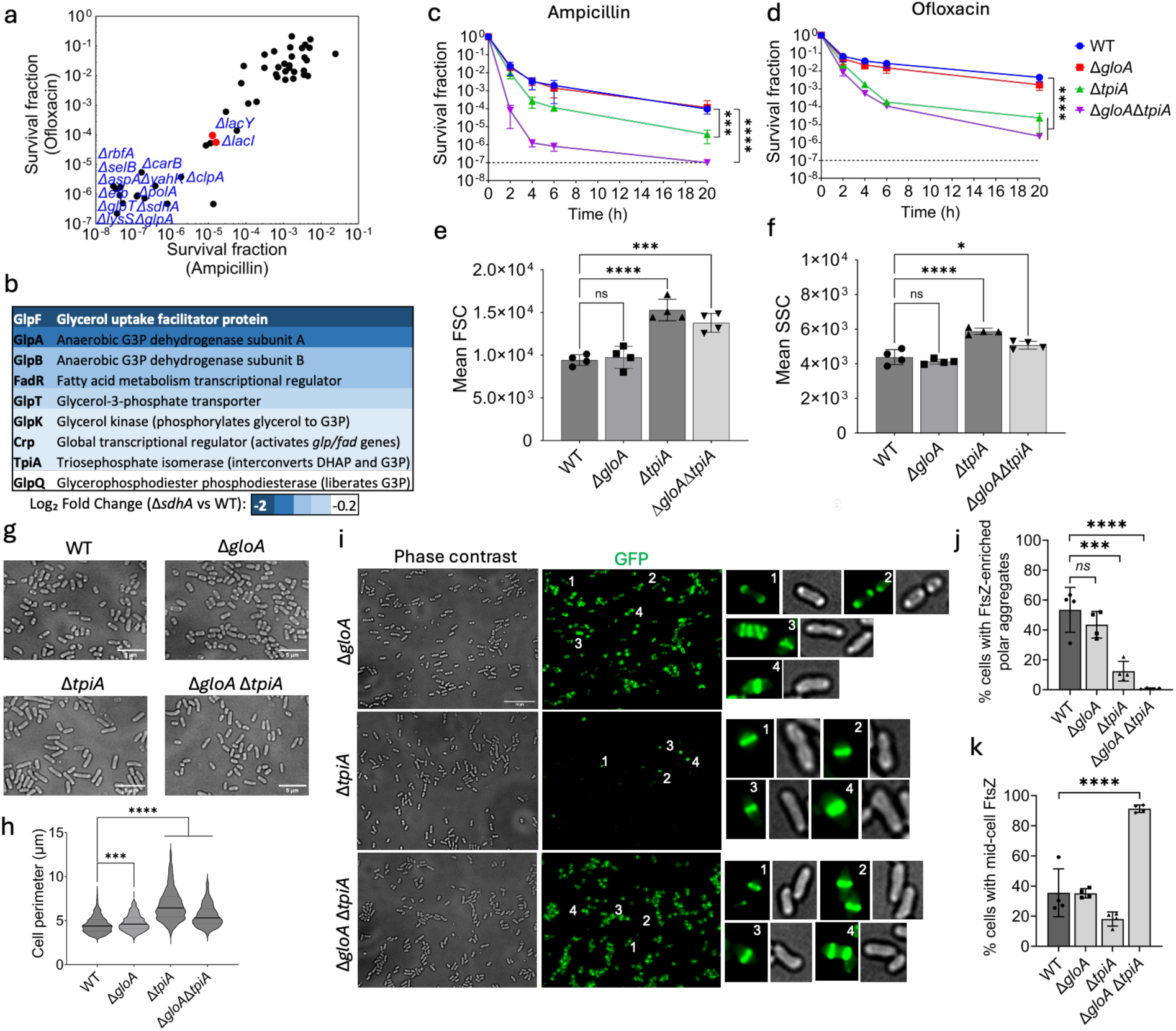
Functional analysis of selected mutants associated with proteomic changes in the Δ*sdhA* strain. (a) High-throughput screening of Keio collection mutants corresponding to genes associated with significantly upregulated or downregulated proteins identified in the proteomics analysis. *lacI* and *lacY* were used as controls. Stationary-phase cells were diluted 100-fold in fresh medium and treated with ampicillin (200 μg/mL) or ofloxacin (5 μg/mL) for 20 hours. Treated cultures were washed, serially diluted, and plated to quantify colony-forming units (CFUs). Survival fraction was defined as the ratio of CFUs after treatment to CFUs measured immediately before antibiotic exposure (N = 1). (b) Network representation of lipid biosynthesis and degradation pathways, highlighting proteins downregulated in the Δ*sdhA* mutant compared to wild type. The color code indicates log₂ fold change (N = 3). (c, d) Kill curves for Δ*gloA*, Δ*tpiA*, and the double knockout Δ*gloA*Δ*tpiA* following treatment with ampicillin (200 μg/mL, panel c) or ofloxacin (5 μg/mL, panel d) (N = 4). (e, f) Forward scatter (e) and side scatter (f) measurements of mutant and wild-type strains by flow cytometry during late stationary phase (N = 4). (g) Phase-contrast micrographs of mutant and wild-type cells. A representative image is shown; similar results were obtained across four biological replicates (N = 4). Scale bar: 5 μm. Cell perimeters were quantified using ImageJ. (h) Quantification of cell perimeters from 1,000 individual cells. (i) Microscope imaging to visualize structural organization of Z-rings and large polar aggregates in *E. coli* WT, Δ*gloA,* Δ*tpiA* and Δ*gloA* Δ*tpiA* strains. Representative phase-contrast and corresponding fluorescence images are shown, with selected cells (numbered) enlarged and displayed side by side to highlight diverse morphologies and structural features. (N = 4). All biological replicates produced similar results. Scale bar, 10 µm. (j) Quantification of the percentage of cells containing large polar aggregates (LPAs) in *E. coli* WT, Δ*gloA,* Δ*tpiA* and Δ*gloA* Δ*tpiA* strains. Each dot represents the mean of a biological replicate, based on analysis of thousands of cells across multiple images (N = 4). (k) Quantification of the percentage of cells containing Z-rings in *E. coli* WT, Δ*gloA,* Δ*tpiA* and Δ*gloA* Δ*tpiA* strains. Each dot represents the mean of a biological replicate, based on analysis of thousands of cells across multiple images (N = 4). All statistical analyses were performed using one-way ANOVA with Dunnett’s multiple comparisons test. For kill curves, ANOVA was performed on the final time points. Statistical significance is indicated as follows: ns, not significant; *P < 0.05; ***P < 0.001; ****P < 0.0001. Data represent the mean ± standard deviation.

To investigate this, we deleted *tpiA*, *gloA*, and both genes in *E. coli* MG1655 and performed selected key assays as described in the previous sections. We specifically focused on TpiA and GloA because they represent metabolic bottlenecks: TpiA maintains carbon flux by interconverting dihydroxyacetone phosphate and glyceraldehyde-3-phosphate, while GloA detoxifies methylglyoxal, a byproduct of elevated glycerol metabolism, feeding into an alternative TCA route (see **Figure 8a**). Although the Δ*tpiA* and the double mutant Δ*gloA*Δ*tpiA* exhibited slower growth compared to the wild type (**Supplementary Figure S7**), flow cytometry–based cell counts confirmed that the total number of cells in late stationary phase was similar across all strains (**Supplementary Figure S8**). Membrane integrity also remained mostly intact in these strains (**Supplementary Figure S9**). While these gene deletions did not drastically alter MICs of *E. coli* (**Supplementary Table S5**), persister levels derived from the stationary phase were significantly reduced in the Δ*tpiA* mutant, with an even more pronounced decrease observed in the double mutant (**Figure 5c, d**). Additionally, since persister cell colonies in the *tpiA* and double mutants grow slowly, assay plates were incubated longer to ensure small colonies had sufficient time to develop and become countable (**Supplementary Figures S10** and **S11**). FSC and SSC analyses revealed that Δ*tpiA* and Δ*gloA*Δ*tpiA* cells were larger and exhibited greater intracellular complexity compared to wild-type cells (**Figure 5e, f**). These findings were validated by microscopy and quantitative measurements of cell dimensions (**Figure 5g, h**). Additionally, Δ*tpiA* and Δ*gloA*Δ*tpiA* mutants displayed reduced levels of LPAs (**Figure 5g**), suggesting improved proteostasis.

To further examine protein aggregation, we quantified FtsZ-GFP localization in Δ*gloA*, Δ*tpiA*, and Δ*gloA*Δ*tpiA* mutants (**Figure 5i)**. Quantitative analysis included 10,622 Δ*gloA*, 6,787 Δ*tpiA*, and 6,150 Δ*gloA*Δ*tpiA* cells. As expected, Δ*gloA* resembled the WT strain, showing no significant differences in LPA or Z-ring formation compared to WT (**Figure 5i, j**). In contrast, both Δ*tpiA* and the double mutant exhibited significantly fewer LPAs (**Figure 5i, j**). Z-ring formation was markedly increased in the Δ*gloA*Δ*tpiA* mutant, but not in Δ*tpiA* alone (**Figure 5i, k**). Non-polar aggregates were rarely observed in the double mutant (**Figure 5i, j, k**). Interestingly, the Δ*tpiA* mutant showed relatively low levels of detectable fluorescence and few Z-rings (which may warrant future independent investigation), but still displayed reduced LPAs compared to WT.

These results (**Figure 5i, j, k**) further reinforce the link between LPAs and persistence, suggesting that polar aggregates support stress tolerance and delayed regrowth, thereby promoting survival under antibiotic challenge. Therefore, we similarly performed time-lapse microscopy of cells exposed to ampicillin. As expected, a substantial fraction of Δ*gloA* cells remained intact after antibiotic treatment, consistent with the presence of LPAs (**Figure 6a**). Even after 5 h of exposure, many Δ*gloA* cells lacking Z-rings but containing LPAs persisted without lysis (**Figure 6a**), similar to WT (**Figure 3a**). On the other hand, Δ*tpiA* and Δ*gloA* Δ*tpiA* mutants were mostly lysed within 3 h, displaying the same morphological sequence observed in the Δ*sdhA* and Δ*sucC* mutants (**Figure 6b,c**).

**Figure 6.**
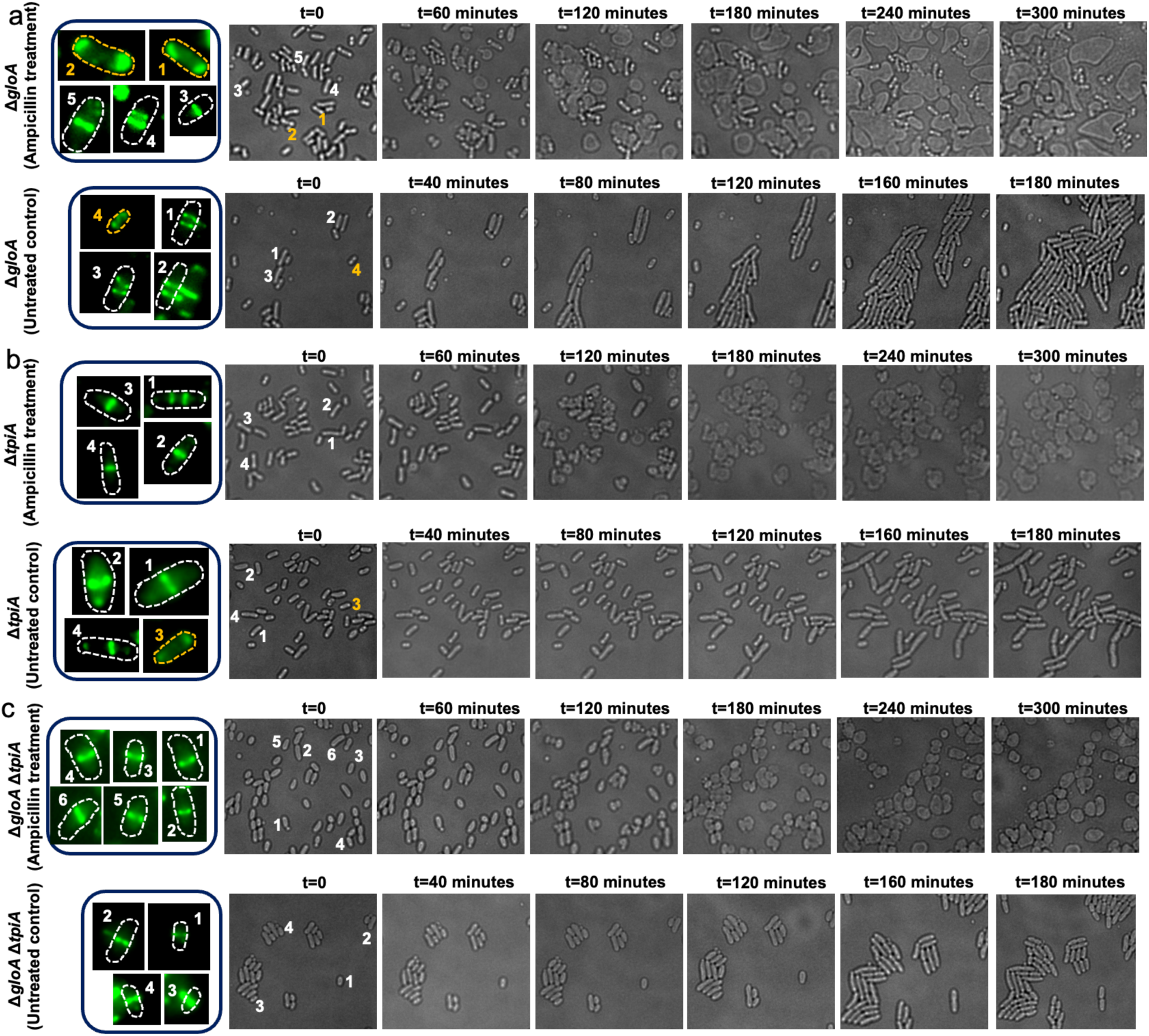
Time-lapse microscopy of (a) Δ*gloA*, (b) Δ*tpiA*, and (c) Δ*gloA*Δ*tpiA* strains under untreated and ampicillin-treated conditions. Each panel contains two rows: ampicillin-treated cells (top) and untreated controls (bottom). In each row, the first column shows a fluorescent image of representative cells at t = 0 h (FtsZ–GFP), with numbered cells outlined by dotted lines to indicate orientation; orange marks non-growing or unlysed cells, and white marks growing or lysed cells. The second column shows the corresponding phase-contrast image at t = 0 h, and subsequent columns show phase-contrast images at the indicated time points. Fluorescent images were acquired only at t = 0 h to visualize Z-rings; repeated blue-laser excitation was avoided due to light sensitivity of the reporter and its potential to perturb growth and lysis dynamics. Images were collected at 37 °C in an on-stage incubator. Cell lysis or growth events can shift cell orientation and position during the time course. A representative image is shown; similar results were obtained across four biological replicates (N = 4).

To validate these observations at the population level and to exclude potential artifacts associated with FtsZ–GFP expression, we performed the same GFP-based lysis control assay described in the previous section using the pUA66-*gfp* reporter. Consistent with the microscopy data, Δ*gloA* cultures retained a substantial population of intact GFP-positive cells following ampicillin treatment, comparable to WT (**Figure 7a, Supplementary Figure S12**). In contrast, Δ*tpiA* and Δ*glo*AΔ*tpiA* cultures exhibited a pronounced loss of GFP signal compared to WT(**Figure 7b,c** and **Supplementary Figure S12**), with the double mutant showing almost no detectable intact cells at the end of the treatment, indicating extensive lysis (**Figure 7c, Supplementary Figure S12**). These orthogonal measurements further support the conclusion that glycerol-dependent metabolic capacity and LPA formation are tightly linked to persistence and antibiotic-induced lysis outcomes.

**Figure 7.**
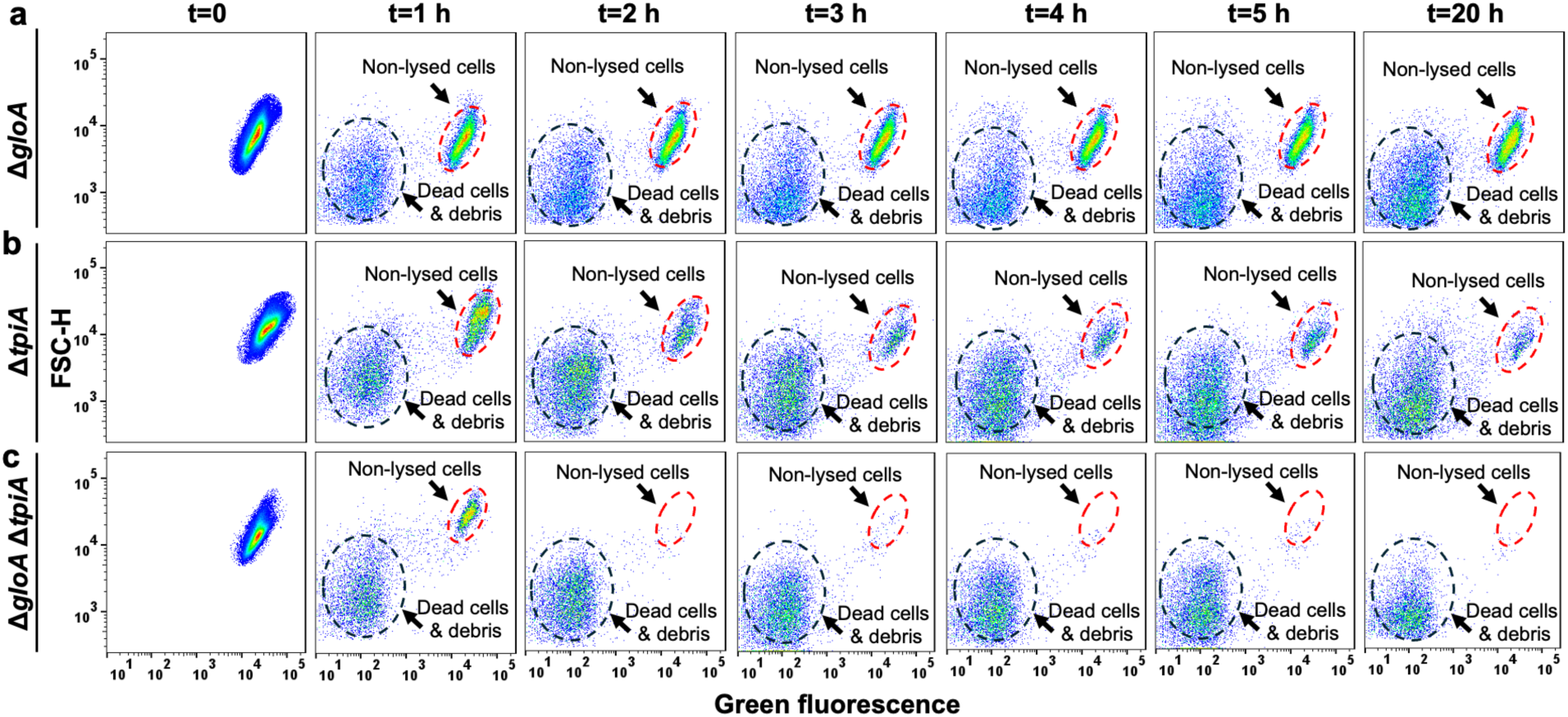
Flow cytometry analysis reveals reduced populations of intact cells in glycerol metabolism mutants. Cells harboring the GFP reporter plasmid pUA66-*gfp* were cultured in the presence of inducer (1 mM IPTG). At the late stationary phase, cultures were diluted 100-fold into fresh medium containing ampicillin and incubated under the same conditions used for microscopy assays. GFP fluorescence of individual cells was monitored by flow cytometry for (a) Δ*gloA*, (b) Δ*tpiA*, and (c) Δ*gloA*Δ*tpiA*. Upon antibiotic-induced lysis, cells lose GFP signal, whereas intact cells retain high GFP fluorescence. Representative flow cytometry profiles are shown; all biological replicates produced similar results (N = 4).

Altogether, these observations are consistent with the phenotypes observed in TCA cycle mutants (**Figures 2** and **3**). The pronounced impact of *tpiA* deletion (**Figure 5c-k**) reflects its central role in maintaining glycolytic flux via DHAP–G3P interconversion, creating a metabolic bottleneck. The further reduction in persistence seen in the double mutant suggests that *gloA*-mediated detoxification of methylglyoxal provides an additional survival advantage under stress.

### Glycerol catabolism sustains PMF in stationary-phase persisters

Given the importance of TCA- and ETC-mediated oxidative phosphorylation in shaping persister phenotypes observed in our results (**Figures 1–3**), and the potential impact of the Δ*gloA*Δ*tpiA* mutant on proton motive force (PMF), a key energy parameter in persister physiology (**Figure 8a**), we directly measured membrane potential using 3,3’-Dipropylthiadicarbocyanine iodide [DiSC_3_(5)]. PMF is the electrochemical potential generated by a difference in proton concentration (ΔpH) and electric charge across a membrane (ΔΨ), mediated by the ETC components such as NADH dehydrogenase, succinate dehydrogenase, cytochromes, and ATP synthase, which transport protons from the TCA cycle products like NADH and FADH₂ across the membrane (**Figure 8a**), and we already shown that genetic perturbation of these key components reduces persistence under the same conditions studied here (thus, these results are not repeated here) ^27^. DiSC_3_(5) is a potentiometric dye that accumulates in polarized membranes, where it self-quenches, resulting in low fluorescence ^69–71^. Upon membrane depolarization, the dye is released into the medium, leading to increased fluorescence (**Supplementary Figure S13**). Conversely, hyperpolarization enhances dye accumulation and quenching, further reducing the fluorescent signal ^69–71^. For these experiments, stationary-phase cells from each strain were incubated with DiSC_3_(5) under identical conditions, and fluorescence was measured after equilibrium was reached. Equilibrium fluorescence reflects the steady-state membrane potential and allows direct comparison between strains. The wild-type strain exhibited higher fluorescence than Δ*gloA*Δ*tpiA* mutants, indicating that the double knockout strain maintains a more hyperpolarized membrane (**Figure 8b**). This shift potentially reflects impaired proton export due to reduced electron transport chain activity and increased intracellular acidification (**Figure 2d**). Importantly, both membrane depolarization and hyperpolarization indicate disruption of PMF homeostasis, reflecting a perturbation of the free energy defined by ΔpH and ΔΨ, which is required for ATP synthesis ^69–71^, and this suggests that the deletions affect membrane energetics in a physiologically substantial way.

**Figure 8.**
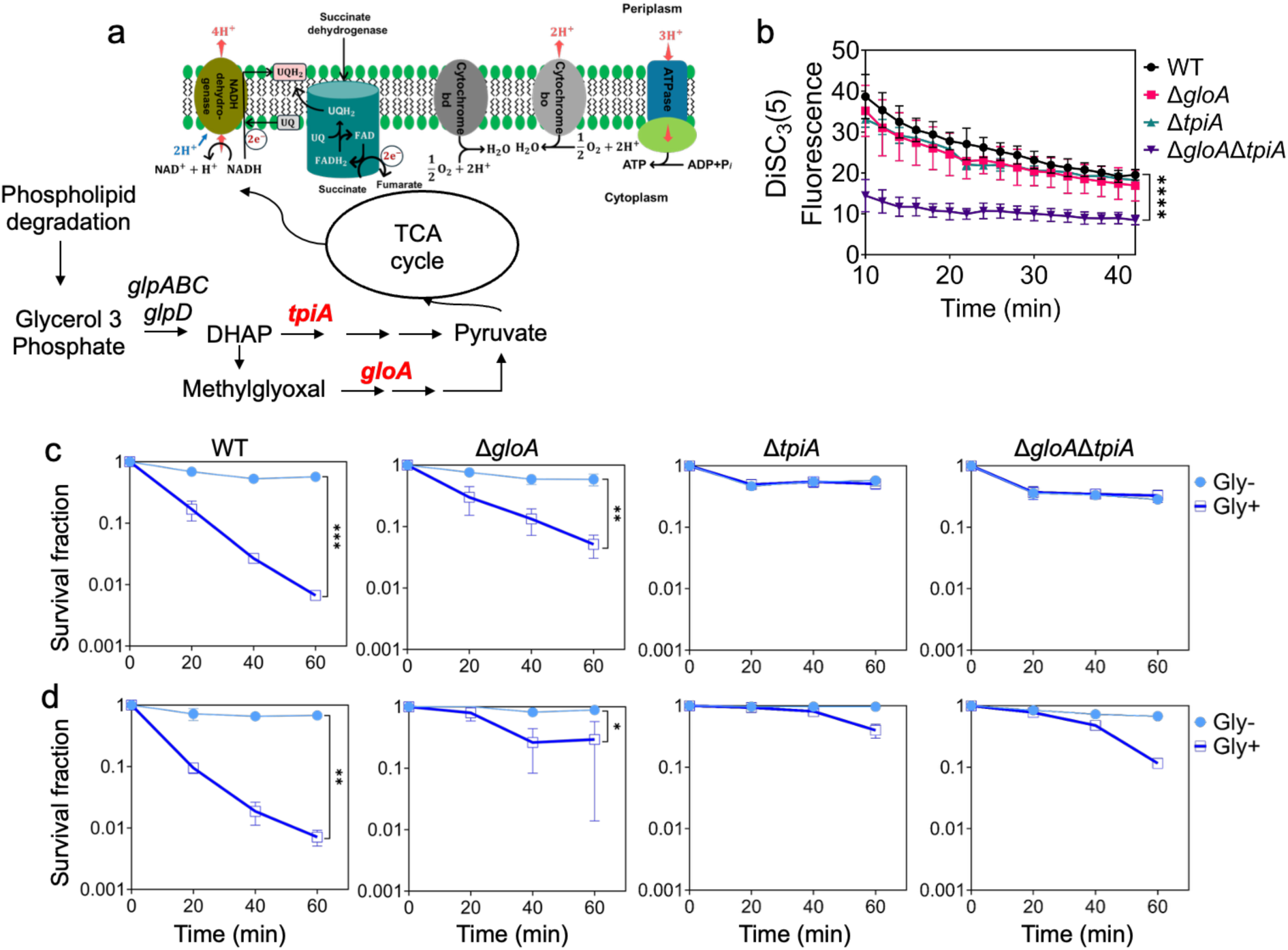
Role of glycerol metabolism in proton motive force (PMF) generation and aminoglycoside (AG) potentiation. (a) Simplified metabolic network illustrating how glycerol metabolism contributes to PMF generation. Together, TpiA and GloA form a metabolic bottleneck for glycerol-derived metabolites feeding into the TCA cycle. (b) PMF measurement using the membrane potential-sensitive dye DiSC₃(5). Equal numbers of cells from each strain at the late stationary phase were incubated with DiSC₃(5), and equilibrium fluorescence was recorded to assess PMF levels. A significant difference in PMF was observed between the WT and the double-knockout strain (N = 4). (c, d) Aminoglycoside potentiation assay in antibiotic-tolerant persister cells. Late stationary-phase cultures were treated with ampicillin (panel c) or ofloxacin (panel d) for 20 hours, then incubated with kanamycin (25 μg/mL) in M9 minimal medium supplemented with either glycerol (Gly+) (60 mM) or deionized water (glycerol negative, Gly-) as a control. Statistical significance testing (see below) was performed for the earlier time points (0, 20, and 40 min) to detect the initial aminoglycoside responses of persister cells (N = 4). Statistical analyses were conducted using simple linear regression models, and F-tests were employed to compare the slopes of the fitted regression lines to assess significant differences among the data sets. Statistical significance is indicated as follows: *P < 0.05; **P < 0.01; ***P < 0.001; ****P < 0.0001. Data represent the mean ± standard deviation.

While DiSC_3_(5) measures PMF at the population level, it does not resolve persister-specific metabolism. To overcome this, we used an aminoglycoside (AG) potentiation assay, which detects PMF-dependent AG uptake triggered by specific metabolites selectively in persister cells ^24^. We previously validated this method as a functional readout of persister metabolism and used it in a high-throughput screen, identifying glycerol and glycerol-3-phosphate as among the most efficiently utilized compounds by persisters ^62^. Although the mechanism was initially unclear to us in that study, revisiting these data alongside our current proteomic and metabolic analyses (**Figures 1-7**) supports the view that stationary-phase persisters retain a selective capacity for energy metabolism, with glycerol catabolism playing a central role in PMF restoration (**Figure 5b**, and **Supplementary Tables S3** and **S4**). To further validate this hypothesis, we applied the AG potentiation assay by resuspending antibiotic-treated wild-type and knockout strains (Δ*tpiA*, Δ*gloA*, and Δ*gloA*Δ*tpiA*) in minimal medium, with or without glycerol, supplemented with an AG (e.g., kanamycin), followed by quantification of CFU levels during the assay ^62^. If persisters actively use glycerol to restore PMF, this rapid metabolic engagement should enhance AG uptake and killing (**Figure 8c, d**). Since ampicillin persisters were around or below the limit of detection (**Figure 5c**), we used high cell density treatment cultures to enrich persister levels (see Materials and Methods); this adjustment did not affect the observed trends (**Supplementary Figure S14**). We specifically focused on AG potentiation responses at early assay time points (**Supplementary Figures S15** and **S16)**, as prolonged incubation could allow persisters to resume growth and obscure their tolerant physiological state. Indeed, persisters arising from late stationary-phase cultures remain in a prolonged drug-tolerant state before growth resumes ^59^, and we have already extensively verified in our previous study that early time-point measurements of AG potentiation assays provide the clearest window into initial metabolic engagement^62^. In the presence of glycerol, both ampicillin and ofloxacin WT persisters showed enhanced AG susceptibility, whereas the Δ*tpiA* and Δ*gloA*Δ*tpiA* mutants did not, and Δ*gloA* showed an intermediate response (**Figure 8c, d,** and **Supplementary Figures S15** and **S16**). To exclude the possibility that the enhanced killing observed in the presence of glycerol reflects persister awakening or growth resumption rather than PMF restoration, we performed parallel control experiments in which antibiotic-treated persisters were incubated with or without glycerol in the absence of aminoglycosides (**Supplementary Figures S17, S18** and **S19**). Over the duration of the AG potentiation assay, no increase in CFU or evidence of cell division was detected in any strain, indicating that glycerol alone does not trigger persister exit or growth under these conditions (**Supplementary Figures S17, S18** and **S19**). Also, we note that in the double mutants, ofloxacin persister levels were slightly reduced during the AG assay (**Figure 8d**). However, this reduction was not specific to AG potentiation, as a similar decrease was observed when these cells were incubated with glycerol in the absence of aminoglycosides (**Supplementary Figure S17**), indicating sustained sensitivity of the double mutants under these conditions even after ending antibiotic treatment. Nevertheless, the increased susceptibility of WT persisters observed upon glycerol addition reflects metabolite-driven PMF reactivation that enables aminoglycoside uptake, rather than persister wake-up or replication. The absence of both GloA and TpiA disrupts this process, establishing it as a critical metabolic bottleneck and further supporting our proposed model linking lipid-derived glycerol catabolism to persister survival.

## Discussion

Our findings reveal a mechanistic link between lipid degradation and glycerol-driven PMF generation in stationary-phase persisters. Studying the stationary phase is particularly relevant, as it reflects the nutrient-limited conditions that bacteria frequently encounter in natural environments, where they are often not actively growing but must still maintain energy homeostasis to survive stress ^17–20^. Proteomic profiling of the TCA cycle mutant, Δ*sdhA*, revealed significant downregulation of enzymes involved in lipid catabolism and glycerol metabolism, suggesting a suppression of endogenous energy-generating pathways that rely on phospholipid turnover. Pathway enrichment analysis reinforced this, highlighting lipid degradation and glycerol utilization as two of the most downregulated metabolic programs. Complementary to these observations, our Keio screen identified multiple genes upstream in lipid and glycerol metabolism whose deletion significantly reduced persistence. These findings align with existing literature; although investigations into these pathways have been sporadic, several studies report that perturbation of specific genes involved in upstream lipid and glycerol metabolism can influence persister levels and/or metabolic activity ^68,72,73^. Notably, AG potentiation assays demonstrate that glycerol and related metabolites are among the most effective carbon sources for sensitizing stationary-phase persisters to aminoglycosides, outperforming other common sugars like glucose, lactose, and fructose ^62^. This effect occurs not through the initiation of new metabolic programs (since phenotypic switching in late stationary phase persisters typically requires extended times) but via retained metabolic capacity that enables rapid PMF restoration upon carbon source availability.

To determine whether reduced persistence in TCA cycle mutants results from changes in oxidative stress, we directly measured DNA oxidation, protein carbonylation, and lipid peroxidation in stationary-phase cultures. Despite observing robust upregulation of oxidative stress response proteins (such as, MsrA/B, SodA, GrxA, and BtuE) in Δ*sdhA*, we found no significant differences in oxidative damage compared to the WT. These findings argue against oxidative macromolecular damage as one of the drivers of persistence in stationary phase. Our results are further supported by prior work under matched experimental conditions, which demonstrated that overexpression of catalases and superoxide dismutases did not affect persister levels, and that anaerobic respiration (absent ROS production) still supported persistence ^32^. Together, these data indicate that while respiration can be necessary for persister formation, it does not act through ROS-induced damage. Although the elevated expression of oxidative stress defense proteins in the TCA cycle mutant seems counterintuitive, it likely reflects a compensatory redox-balancing response to altered or impaired electron flow, rather than an indication of actual oxidative injury. This defense response also contributes to protein homeostasis, potentially explaining the reduced protein aggregation observed in the mutant strains.

In our previous study, we directly tested the role of fatty acid oxidation in stationary-phase persistence by deleting key genes in the β-oxidation pathway, and these deletions had no significant impact on persister formation, indicating that long-chain fatty acid catabolism may not be required for survival under nutrient-limited, stationary-phase conditions ^32^. Our current proteomic analysis highlights the glycerol branch of lipid metabolism, with consistent downregulation of enzymes involved in this pathway in Δ*sdhA*. This metabolic preference makes physiological sense as fatty acid oxidation is highly reducing and energy-rich, but it is also oxygen-intensive and potentially ROS-generating ^74^, conditions that may be unfavorable during stationary phase. In contrast, glycerol may offer a lower-intensity, redox-balanced substrate that supports basal respiration and PMF maintenance. Glycerol can be rapidly converted into central metabolic intermediates ^37^, making it a fast and efficient route to sustain energy homeostasis during prolonged stress.

The marked reduction or absence of LPAs and the increased Z-ring formation in TCA cycle mutants compared to wild-type *E. coli* during stationary phase suggests improved proteostasis and altered stress physiology resulting from metabolic rewiring, and importantly, provides a mechanistic explanation for their reduced persistence. LPAs are known to accumulate in aging, nutrient-depleted cells and have been previously linked to delayed regrowth and increased antibiotic tolerance ^50–53^, making their reduction in TCA mutants particularly notable. Several mechanisms are likely to contribute to this phenotype. First, the lower metabolic activity observed in knockouts (evidenced by reduced RSG fluorescence and downregulation of energy-generating pathways) may reduce proteotoxic stress, limiting the formation of misfolded or damaged proteins^75^. Second, proteomic data revealed upregulation of proteases and redox-balancing enzymes, which enhanced protein quality control and may prevent the accumulation of irreversible aggregates ^76^. Third, elevated ATP levels in most knockouts may support energy-dependent proteostasis pathways, including the activity of chaperones and proteases that maintain a soluble cytoplasmic protein pool ^54,77^. Finally, downregulation of ribosomal proteins and translation factors suggests reduced translational burden, thereby lowering the production of aggregation-prone nascent peptides ^78^.

The decrease in LPA formation also helps explain the unexpected increase in SSC signals in TCA knockouts. In wild-type stationary-phase cells, LPAs localize misfolded proteins into polar foci, potentially removing them from the bulk cytoplasm and reducing cytoplasmic complexity. Additionally, WT cells often undergo significant intracellular degradation under nutrient stress, leading to cellular shrinkage and reduced internal content ^32,79^, both factors that contribute to lower SSC and FSC. This shrinkage is also expected to alter cell membrane architecture, reducing its surface area and potentially releasing phospholipids that can be recycled as internal carbon sources. In contrast, the elevated SSC and FSC in TCA knockouts likely reflects a more intact, granular cytoplasmic architecture (rather than reflecting protein aggregation), and larger cell size. Together, these observations suggest that TCA cycle disruption preserves cellular health during stationary phase and reduces the proteotoxic features associated with persistence. By limiting LPA formation and maintaining protein homeostasis, these cells may avoid the physiological states that promote drug tolerance ^54^, offering mechanistic insight into their reduced persister phenotype.

AG potentiation assays provide a sensitive readout of persister cell metabolism because AGs require active PMF for uptake and lethality ^24,62^. When desired carbon sources are added, persister cells rapidly restores PMF and becomes susceptible to AG killing ^62^. This immediate response (before any significant outgrowth) reflects intrinsic metabolic capability rather than regrowth-driven susceptibility. Importantly, persisters derived from late stationary-phase cultures exhibit extended drug-tolerant states before resuming growth ^59^. Therefore, focusing on early time points allows us to capture the initial metabolic engagement that defines the persister state, avoiding confounding effects of recovery or division. Our data show that glycerol rapidly sensitizes WT persisters to AG, suggesting that these cells either retain latent energy-generating capacity or rapidly activate key metabolic processes such as PMF generation, which itself may require transcriptional and translational activity. This observation supports the central hypothesis of our study that persister cells are not metabolically inert, but instead depend on selective, tightly regulated energy pathways for survival. The ability of persisters to quickly utilize glycerol and restore PMF implies an organized and responsive metabolic infrastructure.

Interestingly, our results also showed a slight reduction in ofloxacin-treated persister levels observed in the double mutants; however, this is not attributable to aminoglycoside potentiation per se. This behavior is not unexpected given the increased antibiotic sensitivity of these mutants. Ofloxacin is known to inflict DNA damage in persister cells as well as in antibiotic-sensitive populations, such that persister survival depends on effective damage repair during the recovery phase ^23^. The increased vulnerability of the double mutants may compromise this repair capacity, leading to reduced apparent persister survival independent of aminoglycoside uptake.

The deletion of key TCA cycle genes actually enhanced survival against aminoglycosides in the exponential phase ^6,47^. Additionally, metabolomic profiling revealed only minor changes in overall metabolic activity and AG uptake in TCA cycle mutants compared to WT during the exponential phase ^47^. The disparity in findings underscores that the metabolic requirements for persistence vary not only by cell type and genetic background but also by the environmental and experimental conditions under which persistence is measured. This discrepancy may be explained by the significant downregulation of ribosomal proteins observed in TCA cycle knockout strains during exponential growth, leading to reduced translation and diminished aminoglycoside efficacy ^47^. Similarly, a recent study showed that ETC (*nuo*) mutants displayed increased aminoglycoside tolerance during exponential growth, a phenotype linked to intracellular acidification and reduced protein synthesis ^49^. In contrast, although our metabolic mutants also exhibited increased cytoplasmic acidification during late stationary phase, these strains showed markedly reduced survival to ampicillin and ofloxacin. These differences suggest that intracellular acidification alone is unlikely to be the primary determinant of persistence in our conditions, and instead may reflect a secondary physiological consequence of disrupted energy metabolism. Together, these observations highlight the adaptive flexibility and redundancy of bacterial survival strategies, which are tightly shaped by physiological state and environmental context.

## Materials and Methods

### Bacterial Strains, Plasmids, Chemicals, Media, and Culture Conditions

*Escherichia coli* K-12 MG1655 wild type (WT) was gift from Dr. Mark P. Brynildsen at Princeton University; plasmid pGFPR01 (the pHluorin reporter) was a gift from Dr. Joan Slonczewski at Kenyon College ^48^; and plasmid pUA66-*ftsZ-gfp* was obtained from our previous study ^59^. Mutant strains of *E. coli* K-12 MG1655 were generated in our previous studies ^27^, and mutants of *E. coli* K-12 BW25113 were obtained from the Keio Collection (Catalog #OEC4988, Horizon Discovery, Lafayette, CO). The *E. coli* K-12 MG1655 Δ*gloA*, Δ*tpiA*, and Δ*gloA*Δ*tpiA* knockout strains were constructed using the Datsenko–Wanner method ^80^. When necessary, the antibiotic resistance cassette was removed via FLP-mediated excision to generate markerless mutants, enabling the generation of double knockout strains ^80^. All strains and primers used to generate and verify the strains in this study are listed in **Supplementary Tables S6** and **S7.** *E. coli* MG1655 strains carrying the low-copy plasmid pUA66-*ftsZ-gfp* were induced with Isopropyl β-D-1-thiogalactopyranoside (IPTG) under the control of a synthetic T5 promoter ^32^. All chemicals were purchased from Fisher Scientific (Atlanta, GA), VWR International (Pittsburgh, PA), or Sigma-Aldrich (St. Louis, MO). Bacterial strains were cultured in liquid Luria–Bertani (LB) medium. LB agar was used for enumerating colony-forming units (CFU). LB broth was prepared by dissolving 5 g of yeast extract, 10 g of tryptone, and 10 g of sodium chloride in 1 L of deionized (DI) water, followed by autoclaving. LB agar was prepared by dissolving 40 g of premixed LB agar in 1 L of DI water. The M9 salt solution was prepared following previously described protocols ^62^. Phosphate-buffered saline (PBS, 1×) was used to wash cells free of antibiotics. Kanamycin (50 μg/mL) was added to overnight and main cultures to maintain plasmid selection. Fluorescent protein expression was induced with 1 mM IPTG. Ampicillin (200 μg/mL) and ofloxacin (5 μg/mL) were used as bactericidal antibiotics in persister assays. The minimum inhibitory concentrations (MICs) of antibiotics for WT and mutant strains were determined using Liofilchem MTS (MIC Test Strips) (Fisher Scientific, Atlanta, GA). Since persisters are known to survive high antibiotic concentrations, we used doses significantly above the MICs (**Supplementary Tables S5**), consistent with the literature. Ampicillin was dissolved in DI water, whereas ofloxacin was dissolved in 0.01 N sodium hydroxide prepared in DI water. Polymyxin B (32 μg/mL), glycerol (600 mM), kanamycin (25 μg/mL), and IPTG (1 M) stock solutions were prepared in DI water. All chemical solutions were sterilized using 0.2 μm VWR syringe filters. Overnight cultures (ONs) were prepared in 14 mL Falcon tubes containing 2 mL of LB medium, inoculated from glycerol stocks (25%) stored at –80°C, and incubated at 37°C with shaking at 250 rpm for 24 h. Main cultures were initiated by diluting ONs 1:1000 in 2 mL of fresh LB medium. Under these conditions, cultures typically reached mid-exponential phase after 3 h and late stationary phase after 24 h. Stationary-phase cells from the main cultures were used for persister, metabolic, and morphological characterization assays, as these cultures reach the stationary phase more uniformly. This consistency results from the controlled initial inoculum size during the main culture setup. In contrast, direct use of ON cultures can lead to variability due to uncontrolled initial cell numbers, often resulting in inconsistent growth phase progression ^32^.

### Cell Growth and Persister Assays

Growth curves of the WT and mutant strains were generated by measuring the optical density at 600 nm (OD_600_) of the main cultures for the indicated time points. At each time point, samples from the main culture and fresh media (used as a blank) were added to a 96-well plate and read at 600 nm using a Varioskan LUX Multimode Microplate Reader (Thermo Fisher, Waltham, MA, USA). Data were collected using SkanIt Software v5.0.

For persister assays, overnight cultures of *E. coli* K-12 (MG1655 WT and BW25113) and their knockout strains were diluted 1:1000 into fresh LB medium. Cultures were prepared in 14 mL Falcon tubes and incubated at 37°C with shaking at 250 rpm. Upon reaching the late stationary phase, the cultures were further diluted 1:100 into fresh LB medium supplemented with antibiotics (ampicillin at 200 μg/mL and ofloxacin at 5 μg/mL). These cultures were then incubated again at 37°C with shaking for 20 hours. After antibiotic treatment, 1 mL from each culture was transferred to a microcentrifuge tube and centrifuged at 13,300 rpm (17,000 × g). The supernatant (950 µL) was discarded, and the cell pellets were washed twice with 1× PBS to reduce residual antibiotic concentrations below the MIC. The washed cells were then resuspended in 100 µL of 1× PBS. A 10 µL aliquot of this suspension was serially diluted in a 96-well round-bottom plate. Subsequently, 10 µL from each dilution was spotted onto LB agar plates. In addition, 90 µL from the most concentrated well was plated to ensure detection of viable cells down to a limit of 1 CFU/mL. Plates were incubated at 37°C for up to 72 hours to allow colony formation. Colony counts were initiated after 16 hours of incubation for fast-growing strains. For strains with slower colony development, 48 hours of incubation was typically sufficient. To generate kill curves, 100 µL samples were collected at the indicated time points during treatment, washed, and plated as described above to quantify changes in CFU over time. For BW25113 mutants, LB medium contained 50 μg/mL kanamycin to prevent potential contamination due to the large number of samples, although each strain was grown in a separate test tube.

### Proteomics Analysis

Proteins were isolated from the *sdhA* knockout and WT strains of *E. coli* K-12 MG1655 at the late stationary phase (24-hour main culture) and sent to UT Health’s Clinical and Translational Proteomics Service Center (Houston, TX) for mass spectrometry analysis. The detailed experimental method is described in our previous study ^27^. Data processing followed the analysis steps outlined by Aguilan et al. ^34^. P-value calculation was performed using a parametric t-test ^34^. The selection of the appropriate t-test type was guided by an F-test, which evaluated whether the replicates for each protein exhibited homoscedasticity (equal variances) or heteroscedasticity (unequal variances) ^34^. The STRING tool (version 12.0) was used to identify significant protein–protein interaction networks among the input proteins ^35^. For protein network and pathway enrichment analysis, we used protein identifiers (accession numbers) for proteins that were upregulated or downregulated by at least 2-fold. STRING-based protein network analysis identified biological pathways that were significantly overrepresented, based on the statistical background of the entire genome and the set of proteins altered in our proteomics analysis. Network edges were defined based on “evidence,” prioritizing different types of interaction evidence to support automated pathway enrichment analysis. Pathway enrichment analysis was performed using multiple functional classification frameworks, including Gene Ontology annotations, KEGG pathways, and UniProt keywords, as previously described ^35^. Enrichment was quantified using a “strength score,” calculated as Log_10_(observed/expected), where higher values indicate stronger enrichment of a given biological pathway. This score reflects the ratio between (i) the number of proteins annotated with a specific term in the observed network and (ii) the expected number in a random network of the same size. Statistical significance was assessed using the False Discovery Rate, with p-values corrected for multiple comparisons within each category using the Benjamini–Hochberg procedure ^35^.

For the volcano plot, we included proteins that were significantly upregulated or downregulated, based on our data analysis following the workflow described by Aguilan et al. ^34^. The threshold for upregulation was set at a fold-change of 2, as indicated by vertical dotted lines on the x-axis. Statistical significance was defined as a p-value ≤0.05, marked by a horizontal dotted line on the y-axis. Color coding was used to illustrate changes in protein levels in the knockout strain compared to wild type: red dots represent upregulated proteins, with corresponding genes listed in the red-outlined box; blue dots represent downregulated proteins, with corresponding genes listed in the blue-outlined box (**Figure 1a**). Genes in both boxes are ordered by the magnitude of fold-change, from highest upregulation to highest downregulation (**Figure 1a**). Gray dots represent proteins that do not meet the significance threshold.

### Flow Cytometry Analysis for FSC and SSC measurements

Main cultures of *E. coli* K-12 MG1655 WT and mutant strains were prepared by diluting 24-hour overnight cultures 1:1000 into 2 mL of LB medium, followed by incubation at 37°C with shaking at 250 rpm until reaching the stationary phase (24 hours). Late stationary-phase cells were then diluted 1:100 in 1 mL of 1× PBS. Prior to flow cytometry analysis, the samples were gently vortexed. Cells were analyzed with a flow cytometer (NovoCyte 3000RYB, Serial #45-1-1612-2663-7, ACEA Biosciences Inc., San Diego, CA) using forward scatter (FSC) and side scatter (SSC) parameters. Samples were run at a flow rate of 14 μL/min, with a sample stream (core) diameter of 7.7 μm. The instrument maintained a constant sheath flow rate of 6.5 mL/min. The core diameter was calculated based on the ratio of the sample flow rate to the sheath flow rate. These conditions were selected to optimize data resolution for measuring *E. coli* cell size. Flow cytometry data were generated using FSC and SSC signals from viable cells. A solvent-only control (without cells) was used to define the background signal and distinguish noise from true cellular events.

### Propidium Iodide (PI) Measurement

Main cultures of *E. coli* K-12 MG1655 WT and mutant strains were prepared by diluting 24-hour overnight cultures 1:1000 into 2 mL of LB medium. Cultures were incubated at 37°C with shaking at 250 rpm and allowed to reach the stationary phase (24 hours). PI dye (Catalog #P1304MP, Thermo Fisher, Waltham, MA) was used to assess membrane integrity. Late stationary-phase cells were diluted 1:100 in 1 mL of 0.85% NaCl buffer. To each sample, 1 μL of 20 mM PI dye was added in flow cytometry tubes. After brief vortexing to ensure uniform mixing, samples were incubated at 37°C in the dark for 10 minutes. Unstained live cells were used to define the gating parameters based on forward and side scatter. Dead cells, treated with 70% ethanol for 30 minutes, served as positive control. Following staining, samples were analyzed by flow cytometry. Cells were excited at 561 nm, and red fluorescence was detected using a 615/20 nm bandpass filter.

### Redox Measurement

Main cultures of *E. coli* K-12 MG1655 WT and mutant strains were prepared by diluting 24-hour overnight cultures 1:1000 into 2 mL of LB medium, followed by incubation at 37°C with shaking at 250 rpm for 24 hours to reach the stationary phase. To measure bacterial metabolic activity, Redox Sensor Green (RSG) dye (Catalog #B34954, Thermo Fisher Scientific, Waltham, MA) was used. Late stationary-phase cells were diluted 1:100 in 1 mL of 0.85% NaCl buffer, and 1 μL of RSG dye was added to each flow cytometry tube. After brief vortexing to ensure uniform mixing, samples were incubated at 37°C in the dark for 10 minutes. Samples were then analyzed by flow cytometry. Cells were excited at 488 nm, and green fluorescence was detected using a 530/30 nm bandpass filter. As a control, cells were treated with 20 μM carbonyl cyanide m-chlorophenyl hydrazone (CCCP) for 5 minutes prior to the addition of RSG dye ^27^.

### Single-cell Analysis of Ampicillin-Induced Cell Lysis with Flow Cytometry

Overnight cultures of *E. coli* MG1655 WT, Δ*sdhA*, Δ*sucC*, Δ*gloA*, Δ*tpiA*, and Δ*gloA* Δ*tpiA* strains harboring the pUA66-*gfp* plasmid were diluted 1,000-fold into 2 mL of fresh LB medium in 14-mL Falcon tubes and grown at 37 °C with shaking for 24 h. At late stationary phase (t = 24 h), cultures were diluted 100-fold into 2 mL of fresh LB medium and immediately treated with ampicillin (200 µg/mL) for up to 20 h. At designated time points (t = 0, 1, 2, 3, 4, 5, and 20 h), treated cells were collected, diluted in 1× PBS, and analyzed by flow cytometry with the same parameters described above. To ensure plasmid maintenance and GFP expression throughout the experiment, cultures were continuously supplemented with kanamycin (50 µg/mL) and IPTG (1 mM), respectively. GFP-positive control samples (cells carrying the pUA66-*gfp* reporter and induced with IPTG) and GFP-negative control samples (cells lacking the reporter plasmid or not induced) were used to define fluorescence thresholds and gate intact cell populations with GFP. Cellular debris and lysed cells were identified based on forward- and side-scatter properties combined with loss of GFP signal. This gating strategy to distinguish intact, lysed, and debris populations has been validated and described in detail in our previous studies ^61,62^. and was applied consistently across all experiments.

### Microscope Imaging for Cell Size Measurement

Main cultures of *E. coli* K-12 MG1655 WT and mutant strains were prepared by diluting 24-hour overnight cultures 1:1000 into 2 mL of LB medium, followed by incubation at 37°C with shaking at 250 rpm until reaching the stationary phase (24 hours). Late stationary-phase cells were then diluted 1:2 with 1× PBS, and 100 μL of the diluted suspension was transferred onto PBS agar pads. The agar pads were prepared by dissolving 1% agarose in 1× PBS, followed by microwave sterilization. The PBS agarose solution was poured over a glass slide (25 × 75 mm), following the method described previously ^59^. A glass coverslip (25 × 75 × 0.17 mm) was placed on top of the agar pad containing the cells. Phase-contrast images were acquired using a fluorescence microscope (Evos FL Auto2, Catalog #AMAFD2000, Thermo Fisher Scientific, Waltham, MA) equipped with a 100× oil immersion objective (Olympus, Catalog #AMEP4733; working distance: 0.3 mm) to assess cellular morphology. ImageJ was used to quantify the cell perimeter in 1,000 cells. To perform this analysis, we first enhanced the contrast of the microscopy images, subtracted the background, set a threshold to remove additional background noise, converted the images to binary masks, and outlined the cell membranes. Finally, we applied filtering conditions to exclude noise and analyze the cells more accurately. Scale bars in the raw images were used as references for cell length measurements.

### LPA and Z-ring Measurements

The method used to image Z-rings in this study was adapted from a previously published approach ^59^. Overnight culture of *E. coli* MG1655 WT, Δ*sdhA*, Δ*sucC*, Δ*gloA*, Δ*tpiA*, Δ*gloA* Δ*tpiA* cells harboring the pUA66-*ftsZ-gfp* plasmid were diluted 1:1000-fold in 2-mL LB broth in 14-mL test tubes and cultured for 24 h with shaking. Kanamycin (50 µg/ml) and IPTG (1 mM) were added in the cultures to maintain the plasmid and to induce *ftsZ-gfp* expression, respectively. At t=24 h, cells were diluted 1:10-fold in PBS (1×). The diluted cells (300 µl) were then transferred to PBS agarose pads, which were dried next to a Bunsen burner flame for ∼20 minutes. The agarose pads were prepared by dissolving agarose (1%) in PBS followed by microwave sterilization. The PBS agarose medium was then poured over a glass slide (25×75 mm), each side of which was enclosed by stacked slides ^59^ to make the pad smooth and sufficiently thick. Once the agar pads containing cells were dried, glass coverslips (25×75×0.1 mm) were placed on the top of the cells. Both phase-contrast and fluorescence (GFP) images were obtained using a fluorescence microscope (Evos FL Auto 2; Catalog # AMAFD2000; Thermo Fisher Scientific) with a 100× (oil) objective (Olympus, Catalog # AMEP4733; working distance, 0.3 mm) to determine cellular morphology and assess expression of the FtsZ-GFP fusion protein. An Evos GFP light cube (Catalog # AMEP4651) was used to acquire GFP images with 470/22-nm excitation and 510/42-nm emission wavelengths. Cell morphology, polar aggregates, and FtsZ ring structures in microscope images were analyzed with ImageJ software. To capture a heterogeneous cell population in each replicate, at least 8 different locations in each agarose pad, leading to a total number of cells ranging from 1000 to 2500 (x4 biological replicates), were selected and monitored with the use of the Evos FL Auto 2 imaging system. The brightness of the phase-contrast images and the color of the fluorescent images were adjusted with “brightness/contrast,” “color balance,” and “sharpen” options. The polar aggregates and Z-rings within a bacterium was enumerated manually using the fluorescent images.

### Time-lapse microscope imaging

Main cultures of *E. coli* K-12 MG1655 WT and mutant strains were prepared by diluting 24 h overnight cultures 1:1000 into 2 mL of LB medium and incubating at 37 °C with shaking (250 rpm) until stationary phase (24 h). Late stationary-phase cells were then diluted 1:5 in fresh LB medium, and 100 μL of the suspension was transferred onto LB agarose pads. Agarose pads were prepared by dissolving 1% agarose in LB medium, sterilizing the solution by microwaving, supplementing with ampicillin, and pouring it onto glass slides (25 × 75 mm). To maintain the pUA66*-ftsZ-gfp* plasmid and induce expression, 50 μg/mL kanamycin and 1 mM IPTG were added, respectively. For ampicillin treatment, a final concentration of 200 μg/mL was used in agar pads, consistent with the persister assay. Untreated pads served as controls. Time-lapse imaging was performed using an on-stage incubator to maintain 37 °C, air, and humidity. Phase-contrast and fluorescence (GFP) images were captured with an Evos FL Auto 2 fluorescence microscope equipped with a 100× oil immersion objective (specifics provided above). Cellular morphology and FtsZ-GFP expression were analyzed. GFP fluorescence was detected using an Evos GFP light cube with 470/22-nm excitation and 510/42-nm emission filters.

### ATP Measurement

Main cultures of *E. coli* K-12 MG1655 WT and mutant strains were prepared by diluting 24-hour overnight cultures 1:1000 into 2 mL of LB medium in 14 mL Falcon tubes, followed by incubation at 37°C with shaking at 250 rpm for 24 hours to reach the stationary phase. To determine cell numbers, 10 μL of the main culture at the stationary phase was analyzed by flow cytometry, as described above. Simultaneously, 100 μL of the main culture was mixed with 100 μL of ATP detection reagent from the BacTiter-Glo™ Microbial Cell Viability Assay (Catalog #G8230, Promega, Wisconsin, USA) in a black, flat-bottom 96-well plate. The mixture was incubated for 5 minutes at 37°C on the same shaker. Luminescence was measured using the Varioskan LUX Multimode Microplate Reader (Thermo Fisher, Waltham, MA, USA). Data were collected using SkanIt Software v5.0, and ATP levels were calculated based on a standard curve generated from known ATP concentrations and normalized to cell numbers.

### pH Measurement

A standard curve for intracellular pH as a function of fluorescence ratio (410/470) in *E. coli* K-12 MG1655 was generated using the previously described steps ^48,49^. Cells carrying the pGFPR01 plasmid encoding pHluorin were induced with 0.2% L-arabinose in the main culture to express the pH-sensitive fluorescent protein. The plasmid was maintained with 100 μg/mL ampicillin. To create pH buffers necessary for the standard curve, 2-(N-morpholino) ethanesulfonic acid (MES) and 3-(N-morpholino) propanesulfonic acid (MOPS) solutions were prepared at 50 mM concentrations, adjusted to pH values ranging from 5.0 to 10.0 in 0.5-unit increments. For the standard curve, first, transmembrane pH collapsed by adding 40 mM potassium benzoate and 40 mM methylamine hydrochloride to the stationary phase cultures, equalizing the difference between the external and internal pH. Then, culture samples were collected, washed to remove the chemicals, and normalized to an OD_600_ of 1.0 in 0.5 mL of each pH buffer. A 150 μL aliquot from each suspension was transferred into a 96-well plate, with LB used as the blank. The fluorescence ratio (410/470) was measured using a plate reader, and the generated standard curve was used to obtain a Boltzmann sigmoid best-fit equation ^49^:

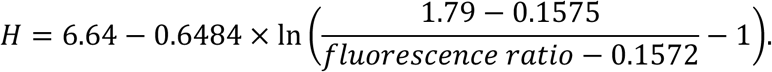

For pH measurements in experimental samples, main cultures of *E. coli* K-12 MG1655 WT and mutant strains carrying the pGFPR01 plasmid (maintained with 100 μg/mL ampicillin and induced with 0.2% L-arabinose) were prepared by diluting 24-hour overnight cultures 1:1000 into 2 mL of LB medium. Cultures were incubated in 14 mL Falcon tubes at 37°C with shaking at 250 rpm and grown to the stationary phase (24 hours). Then, OD_600_ of the cultures was normalized to 1.0 in fresh LB, and 150 μL from these normalized cultures was transferred to a 96-well plate (LB as blank), and the fluorescence ratio was measured with a plate reader. This ratio was then used to determine pH based on the standard curve equation above.

### DNA Damage Assay

Main cultures of *E. coli* K-12 MG1655 WT and mutant strains were prepared by diluting 24-hour overnight cultures 1:1000 into 2 mL of LB medium, followed by incubation at 37°C with shaking at 250 rpm for 24 hours to reach the stationary phase. Cell densities were normalized based on the OD_600_ of WT and mutant strains at 2.5, in 2 mL samples, which were then used for total DNA extraction. DNA extraction and quantification were performed using the DNeasy Blood & Tissue Kit (Catalog #69504, Qiagen, Germantown, MD, USA) according to the manufacturer’s instructions. DNA at a concentration of 25 ng/µL in a total volume of 45 µL was used for the DNA damage assay. To the extracted DNA, 5 µL of 10× Nuclease P1 Reaction Buffer (Catalog #M0660, New England BioLabs Inc., Ipswich, MA, USA) and 0.1 µL of Nuclease P1 were added. The pH of the solution was adjusted using 1 M Tris buffer, followed by the addition of 1 unit of alkaline phosphatase (*E. coli* C75) (Catalog #2120A, Takara Bio Inc., Kusatsu, Shiga, Japan). The mixture was incubated at 37°C for 30 minutes, then the reaction was inactivated by heating at 75°C for 10 minutes. DNA damage levels were measured using the DNA/RNA Oxidative Damage (High Sensitivity) ELISA Kit (Catalog #589320, Cayman Chemical, Ann Arbor, MI, USA) according to the manufacturer’s protocol. This assay measures DNA/RNA damage caused by oxidative processes. Specifically, it detects oxidatively damaged guanine species, including 8-hydroxyguanosine, 8-hydroxy-2’-deoxyguanosine, and 8-hydroxyguanosine. The assay is an ELISA that utilizes AChE (8-OH-dG–acetylcholinesterase conjugate) technology. Equal amounts of extracted DNA from each sample are digested, and the lysates are incubated with AChE to compete for binding to antibodies specific to DNA/RNA oxidative damage. These antibody-bound complexes then attach to goat polyclonal anti-mouse IgG coated on the bottom of the 96-well plate provided by the vendor. After washing away unbound substances, the bound complexes are quantified using a reagent supplied by the vendor, which catalyzes an enzymatic reaction producing a distinct yellow color measured at 412 nm. The intensity of this color is proportional to the amount of AChE complexed with the antibodies and inversely proportional to the level of oxidative damage in the DNA/RNA samples. A standard curve for the DNA/RNA oxidative damage ELISA was generated using the assay’s provided standard solution.

### Protein Damage Assay

Main cultures of *E. coli* K-12 MG1655 WT and mutant strains were prepared by diluting 24-hour overnight cultures 1:1000 into 2 mL of LB medium, followed by incubation at 37°C with shaking at 250 rpm for 24 hours to reach the stationary phase. Cell densities were normalized based on an OD_600_ of 2.5 for both WT and mutant strains, and 2 mL of these cells were washed twice with cold 1× PBS while kept on ice throughout the process. Centrifugation was performed at 4°C, 13,000 rpm for 3 minutes. The resulting pellets were lysed in 300 µL of NEBExpress *E. coli* Lysis Reagent (Catalog #P8116S, New England BioLabs Inc., Ipswich, MA, USA) at room temperature for 30 minutes. The lysate was then centrifuged at 16,600 × g for 10 minutes, and 250 µL of the supernatant was collected for the assay. Total protein concentration was determined using the Bicinchoninic Acid (BCA) assay (Catalog# 23225, Thermo Fisher Scientific, Waltham, MA). Twenty-five µL of each cell lysate sample and DI water as a blank were loaded into a 96-well plate. Then, 200 µL of BCA working solution (prepared at a 50:1 ratio of Reagent A to B) was added to each well and mixed thoroughly on a plate shaker for 30 seconds. The plate was incubated at 37°C for 30 minutes and then cooled for 5 minutes at room temperature, protected from light. Absorbance was measured at 562 nm using a plate reader. Protein concentrations were calculated using a standard curve generated from BCA standards. Protein carbonyl levels in control and mutant strains were measured using the Protein Carbonyl Colorimetric Assay Kit (Catalog #10005020, Cayman Chemical, Ann Arbor, MI, USA) according to the manufacturer’s instructions and the detailed analysis protocol. This assay measures protein oxidation by quantifying carbonyl protein content, a common marker of oxidative damage. Carbonyl groups form when iron and copper cations bind to proteins and are subsequently attacked by reactive oxygen species, which oxidize the side-chain amine groups of certain amino acids into carbonyls. The lysed samples are then incubated with 2,4-dinitrophenylhydrazine (DNPH). DNPH reacts with the carbonyl groups to form Schiff bases, producing hydrazones that can be analyzed spectrophotometrically at 360 to 385 nm.

### Lipid Damage Assay

Main cultures of *E. coli* K-12 MG1655 WT and mutant strains were prepared by diluting 24-hour overnight cultures 1:1000 into 2 mL of LB medium, followed by incubation at 37°C with shaking at 250 rpm for 24 hours to reach the stationary phase. Cell densities were normalized based on an OD_600_ of 2.5 for both WT and mutant strains. 1 mL of late stationary phase cells was washed twice with cold 1× PBS while kept on ice throughout the process. Centrifugation was performed at 4°C, 13,000 rpm for 3 minutes. After the final centrifugation, all the supernatant was removed except for 100 µL, which was used to resuspend the cell pellets. Cell density was measured by flow cytometry using 10 µL of the adjusted culture diluted in 990 µL of 1× PBS. The remaining cell culture was treated with a 100-fold excess of butylated hydroxytoluene solution to prevent further oxidation and then lysed using the SDS cell lysis buffer provided in the OxiSelect™ TBARS Assay Kit (Catalog #STA-330, Cell Biolabs, Inc., San Diego, CA, USA). The mixture was incubated at room temperature for 5 minutes, followed by the addition of 250 µL of thiobarbituric acid reagent. The mixture was then incubated for 60 minutes at 95°C and subsequently cooled in an ice bath for 5 minutes. After centrifugation at 13,000 rpm for 15 minutes, the supernatant was collected for further analysis. 200 µL of the supernatant were added to a 96-well plate and read at 532 nm using a Varioskan LUX Multimode Microplate Reader (Thermo Fisher, Waltham, MA, USA). Data were collected using SkanIt Software V 5.0. A standard curve for malondialdehyde (MDA) was generated using the assay’s provided standard solution. This assay measures lipid peroxidation by detecting its end-products, such as MDA, which is stable and a widely accepted marker of oxidative stress. The lysed samples are incubated with thiobarbituric acid reagent, where MDA forms a complex with thiobarbituric acid reactive substances (TBARS). This complex can be measured calorimetrically at 532 nm or fluorometrically with excitation at 540 nm and emission at 590 nm.

### DiSC_3_(5) Assay

The DiSC_3_(5) assay was performed as previously described ^69–71^. Main cultures of *E. coli* K-12 MG1655 WT and mutant strains were prepared by diluting 24-hour overnight cultures 1:1000 into 2 mL of LB medium, followed by incubation at 37°C with shaking at 250 rpm until reaching stationary phase at 24 hours. Late stationary-phase cells were collected, normalized to an OD_600_ of 0.1, and washed twice with buffer containing 5 mM HEPES (N-[2-Hydroxyethyl]piperazine-N-[2-ethanesulfonic] hemisodium salt) and 20 mM glucose. The washed cells were stained with 1 μM DiSC_3_(5) diluted 1:1000 and incubated in the dark. Fluorescence levels were measured using a plate reader at 620 nm excitation and 670 nm emission wavelengths every 2 minutes for up to 42 minutes. After 20 minutes, when fluorescence levels reached equilibrium, control samples were treated with polymyxin B (32 μg/mL), a membrane disruptor that binds lipopolysaccharide and induces conformational changes in the membrane, and fluorescence was remeasured for another 20 minutes. Polymyxin B treatment served to verify the functionality of the dye.

### Aminoglycoside Potentiation Assay

The AG potentiation assay was performed as previously described ^62^, with some modifications. The main culture was diluted 10-fold into 25 mL of fresh media in 250 mL baffled flasks and treated with ampicillin (200 μg/mL) and ofloxacin (5 μg/mL) for 20 hours. Persisters were collected from the 25 mL culture, washed twice with M9 salt medium, suspended in 1 mL (cells concentrated), and divided into equal cell numbers into two test tubes containing either DI water (control) or 60 mM glycerol. Both sets were subsequently treated with kanamycin (25 μg/mL). Every 20 minutes, 100 μL samples were taken, washed twice with 1× PBS, serially diluted in a 96-well plate, and 10 μL from each dilution was plated onto LB agar plates. Colony-forming units were quantified after incubation at 37°C until visible colonies appeared, for up to 72 hours.

### Statistical Analysis

Each experiment included at least three biological replicates unless otherwise stated. Data points in the figures represent the mean ± standard deviation. For microscopy analyses, representative images from both phase-contrast and fluorescence microscopy are shown. Statistical analyses were performed using one-way ANOVA with Dunnett’s multiple comparisons test where applicable. For time-dependent responses, simple linear regression models were used, and F-tests were applied to compare the slopes of the regression lines. Thresholds for significance were set as follows: *P < 0.05, **P < 0.01, ***P < 0.001, ****P < 0.0001. GraphPad Prism and FlowJo were used to generate figures and perform statistical analyses.

## Acknowledgment

The authors would like to thank the members of Orman Lab for their help. This study was supported by NSF CAREER 2044375, NSF 2326142, and NIH/NIAID R01-AI143643.

Proteomics experiments were conducted at the Mass Spectrometry Laboratory of Dr. Chengzhi Cai at the University of Houston, with the associated service fees.

## Contributions

H.N., S.G.M, and M.A.O. conceived and designed the study. H.N. and S.G.M. performed the experiments. H.N., S.G.M., and M.A.O. analyzed the data and wrote the paper. All authors have read and approved the manuscript.

## Data availability

All data supporting this manuscript are available in the main text or the supplementary file. The complete proteomics dataset is provided as “Supplementary Proteomics Data.”

## Movie

**Movie 1. Time-lapse microscopy of *E. coli* cells during ampicillin treatment.**

Late stationary-phase cells were immobilized on LB agarose pads containing ampicillin (200 µg/mL) and imaged at 37 °C in an on-stage incubator. Lysis can be a rapid and dynamic process, with cells collapsing into spherical spheroplasts or protoplasts before rupturing and releasing their intracellular contents (indicated by arrows). We note that ampicillin induced the same lysis dynamics in all *E. coli* strains tested.

