## Supplementary figures and tables for "Glycerol-Driven Energy and Proteostasis Underpin Antibiotic Tolerance in *Escherichia coli*"

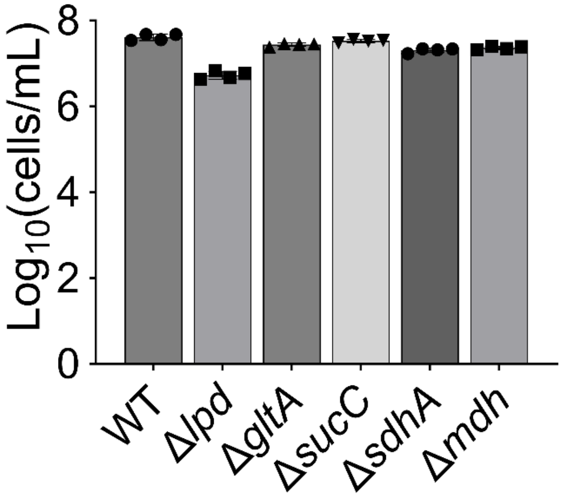


**Fig. S1. Flow cytometry-based cell quantification.** *Escherichia coli* wild-type (WT) and mutant strains (Δ*lpd*, Δ*gltA*, Δ*sucC*, Δ*sdhA*, and Δ*mdh*) at late stationary phase (24 h) were diluted in 1× PBS and analyzed by flow cytometry to quantify cell counts. The number of biological replicates, N=4. Data represent the mean ± standard deviation.

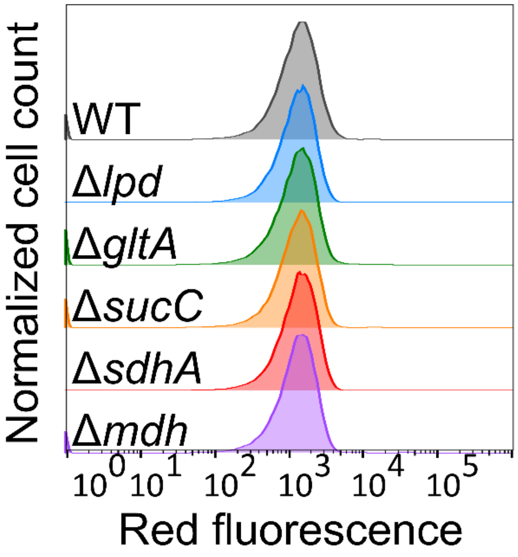


**Fig. S2. Propidium iodide staining of cells at late stationary phase.** *E. coli* WT and mutant strains (Δ*lpd*, Δ*gltA*, Δ*sucC*, Δ*sdhA*, and Δ*mdh*) at late stationary phase (24 h) were collected, diluted in sterile 0.85% NaCl buffer solution, and then stained with 20 μM propidium iodide dye. Stained cells were analyzed by flow cytometry. A minimum of four biological replicates were conducted, with each displaying consistent trends. The flow diagram shown is representative of these results.

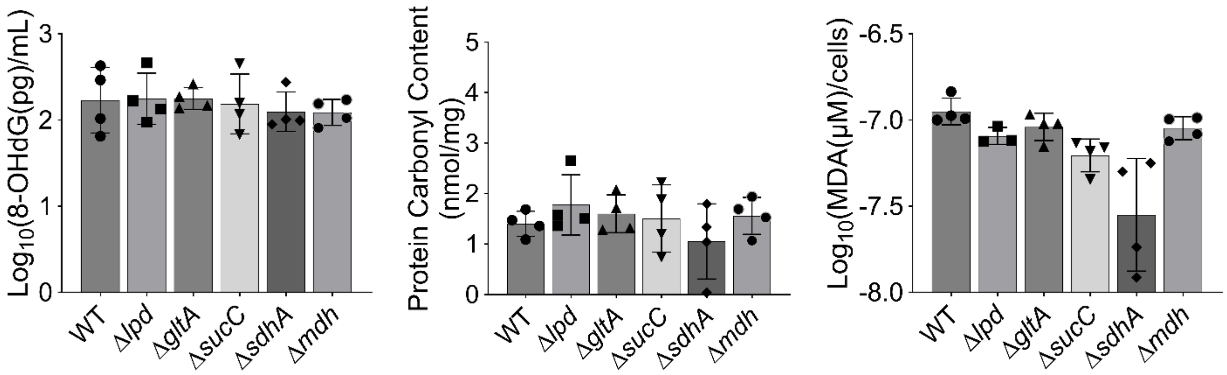


**Fig. S3. Assessment of oxidative stress using three biomarkers: DNA oxidation, protein carbonylation, and lipid peroxidation.** Cells from the indicated strains at late stationary phase (24 h) were collected to assess DNA oxidation, protein carbonylation, and lipid peroxidation. These parameters were measured using commercial kits following the manufacturer's protocols (see **Materials and Methods**). For each condition and strain, identical cell densities were used. Protein carbonyl content and lipid damage (MDA) levels were further normalized to total protein concentration and cell number, respectively. DNA damage levels (8-OHdG) are reported as absolute concentrations, as the same amount of DNA was used for all conditions and strains. N=4. Data represent the mean ± standard deviation.

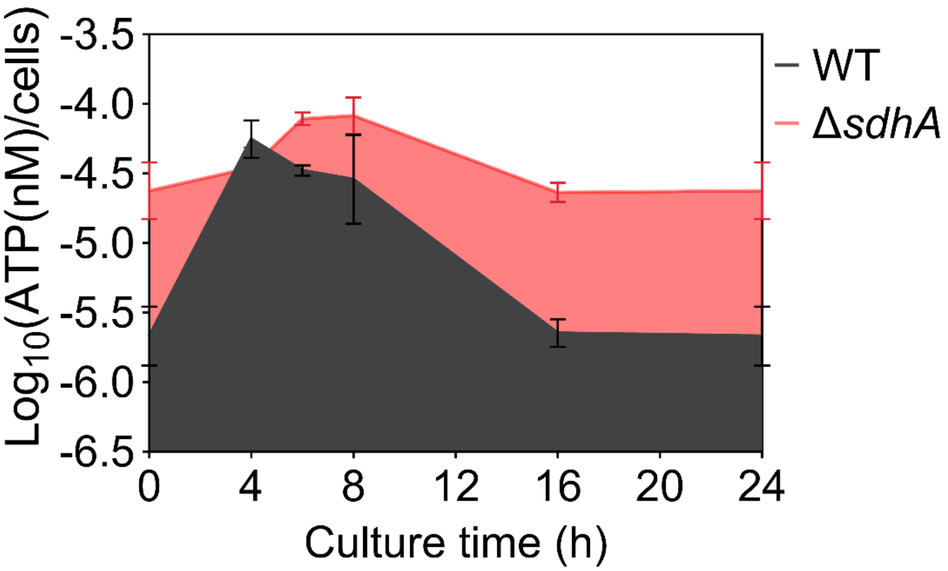


**Fig. S4. Quantification of intracellular ATP levels in the cells.** Overnight stationary phase cultures of *E. coli* WT and Δ*sdhA* strains were diluted 1:1000-fold into fresh medium and incubated for 24 hours. At specified time points, cells were collected, and intracellular ATP levels were measured using the BacTiter-Glo™ Microbial Cell Viability Assay Kit, following the manufacturer's instructions. To normalize ATP concentrations per cell, cell counts were simultaneously determined using flow cytometry. N=4. Data represent the mean ± standard deviation.

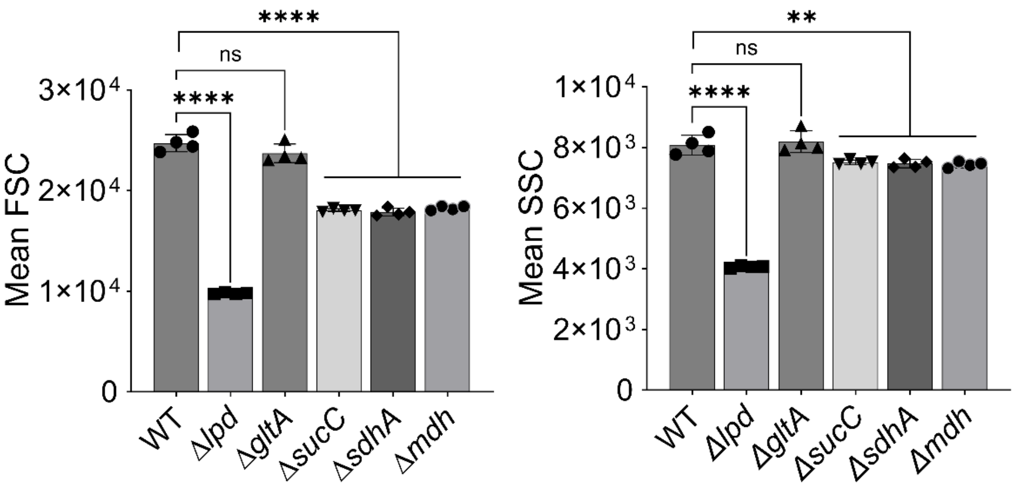


**Fig. S5. Morphological characterization of TCA cycle knockout strains during the exponential phase.** Overnight cultures of *E. coli* WT and mutant strains (Δ*lpd*, Δ*gltA*, Δ*sucC*, Δ*sdhA*, and Δ*mdh*) were diluted 1:1000-fold in fresh LB media and grown to exponential phase (t=3 h). Then, cells were collected and diluted in 1× PBS and analyzed by flow cytometry to measure forward scatter (FSC) and side scatter (SSC) at the single-cell level. N=4. Statistical analysis was performed between the WT and single mutants using one-way ANOVA with Dunnett's post-test. *P<0.05, **P < 0.01, ***P < 0.001, and ****P < 0.0001. Data represent the mean ± standard deviation.

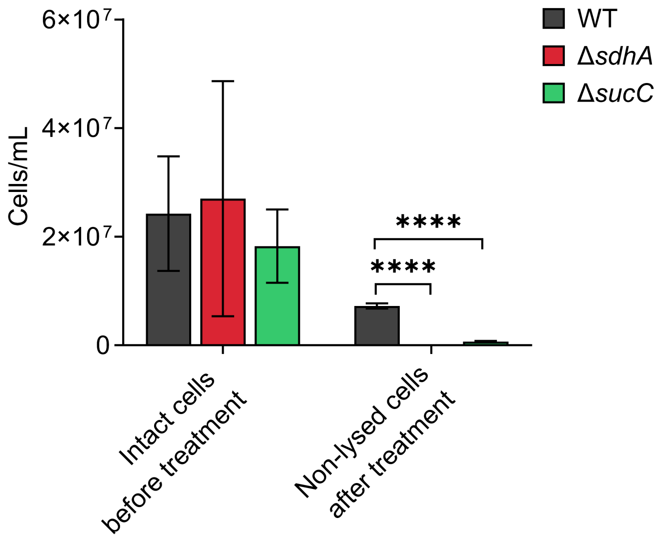


**Fig. S6. Flow cytometry quantification of intact cells after ampicillin treatment.** The fraction of intact (GFP-positive) cells was measured by flow cytometry before and after 20 h of ampicillin treatment in WT, Δ*sdhA*, and Δ*sucC* strains. N = 4. Statistical analysis was performed using two-way ANOVA with Dunnett's multiple comparisons test; ****$P < 0.0001$. Data represent mean ± standard deviation from four independent biological replicates.

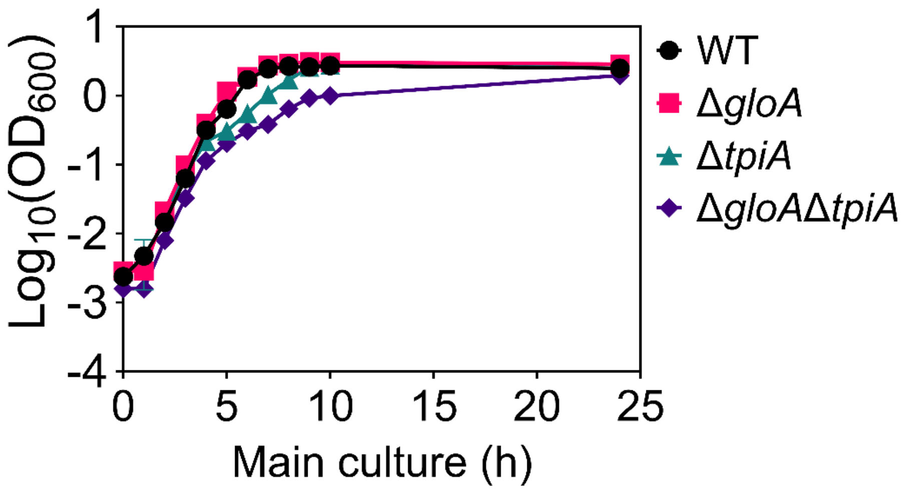


**Fig. S7. Growth curves of WT, Δ*gloA*, Δ*tpiA*, and Δ*gloA* Δ*tpiA* strains.** Overnight cultures of *E. coli* WT and mutant strains (Δ*gloA*, Δ*tpiA*, and the double mutant Δ*gloA* Δ*tpiA*) were diluted 1:1000-fold into fresh LB medium and incubated for 24 hours at 37 °C with shaking. At designated time points, cells were collected, and optical density at 600 nm ($OD_{600}$) was measured using a plate reader. N=4. Data represent the mean ± standard deviation.

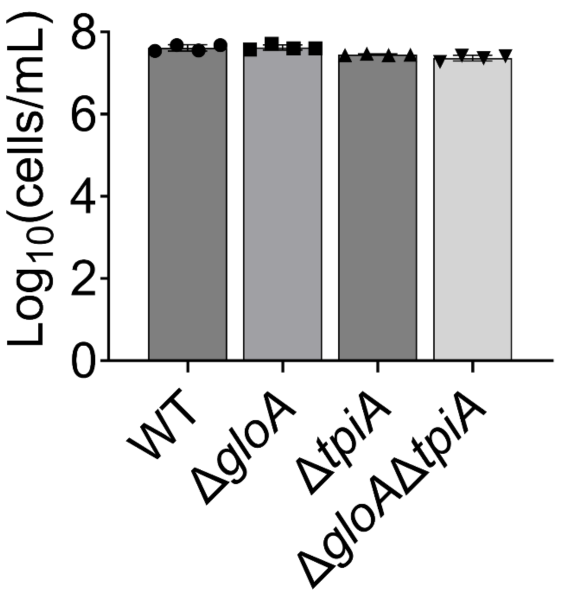


**Fig. S8. Flow cytometry-based cell quantification.** *E. coli* WT and mutant strains (Δ*gloA*, Δ*tpiA*, and the double mutant Δ*gloA* Δ*tpiA*) at late stationary phase (24 h) were diluted in 1× PBS and analyzed by flow cytometry to quantify cell counts. N=4. Data represent the mean ± standard deviation.

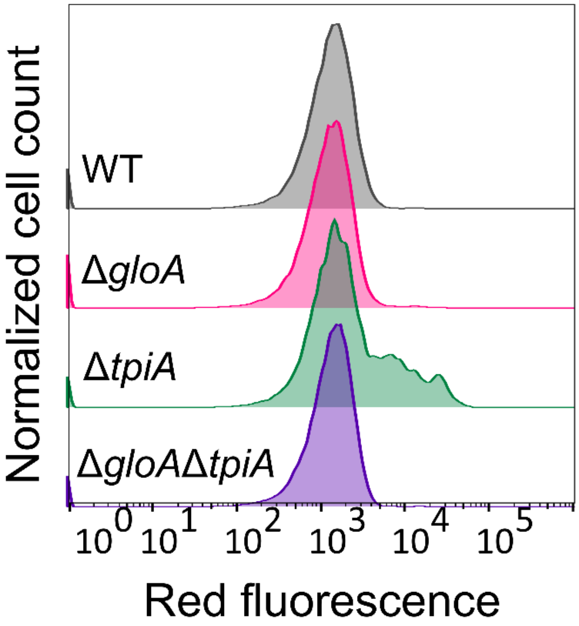


**Fig. S9. Propidium iodide staining of cells at late stationary phase.** *E. coli* WT and mutant strains (Δ*gloA*, Δ*tpiA*, and the double mutant Δ*gloA* Δ*tpiA*) at late stationary phase (24 h) were collected, diluted in sterile 0.85% NaCl buffer solution, and then stained with 20 μM propidium iodide dye. Stained cells were analyzed by flow cytometry. A minimum of four biological replicates were conducted, with each displaying consistent trends. The flow diagram shown is representative of these results.

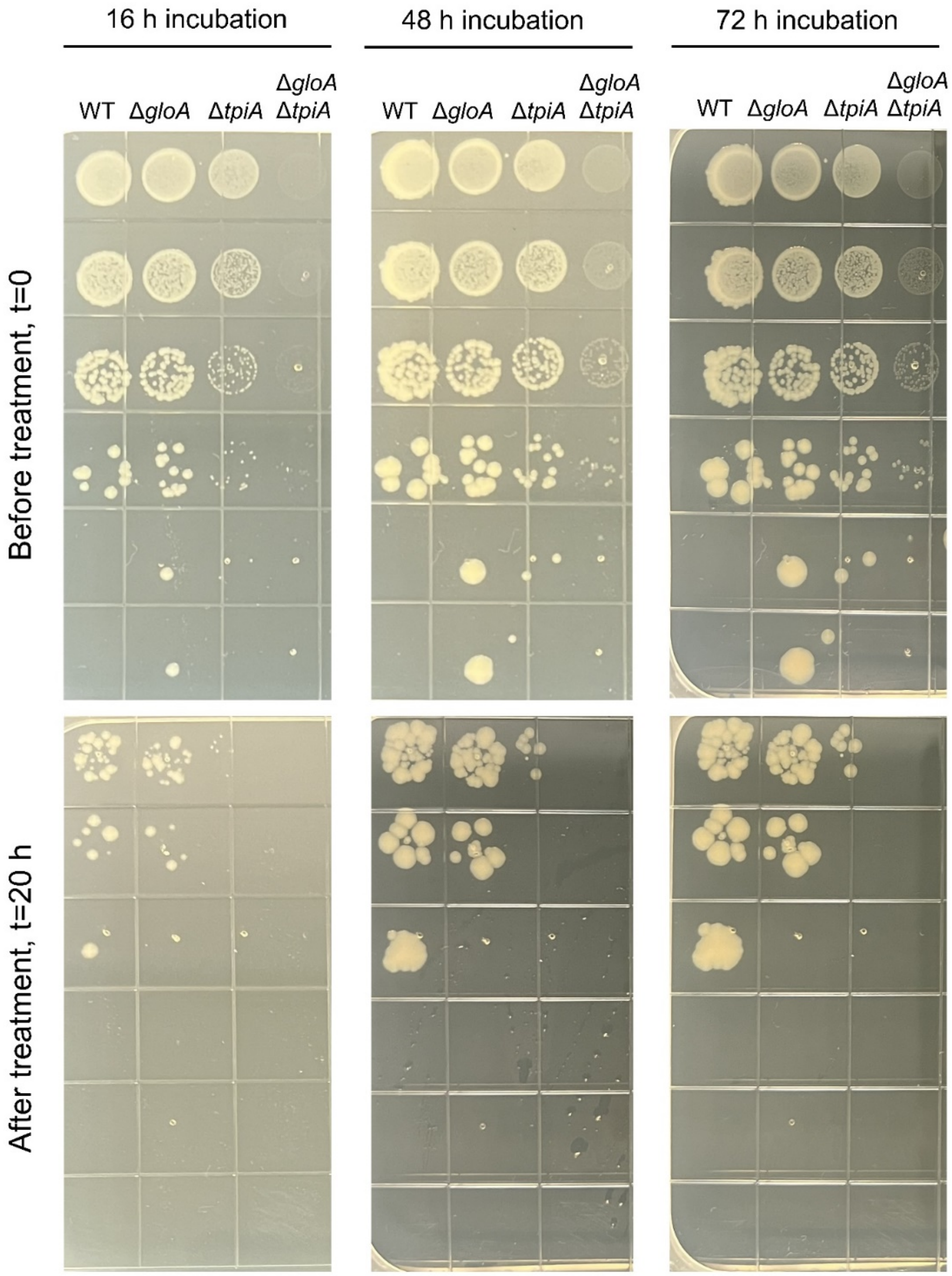


**Fig. S10. Colony formation of WT and mutant strains visualized on agar plates.** Late-stationary-phase (24 h) cultures of *E. coli* WT, Δ*gloA*, Δ*tpiA*, and the double mutant Δ*gloA* Δ*tpiA* were diluted 1:100-fold into fresh medium and exposed to ampicillin (200 µg/ml) for 20 h. After the treatment, cells were harvested, washed with 1× PBS, and plated on LB agar to count colony formation units (CFU). Plates were incubated for 72 h to allow small colonies to develop fully, and images were captured at specified time points. N=4.

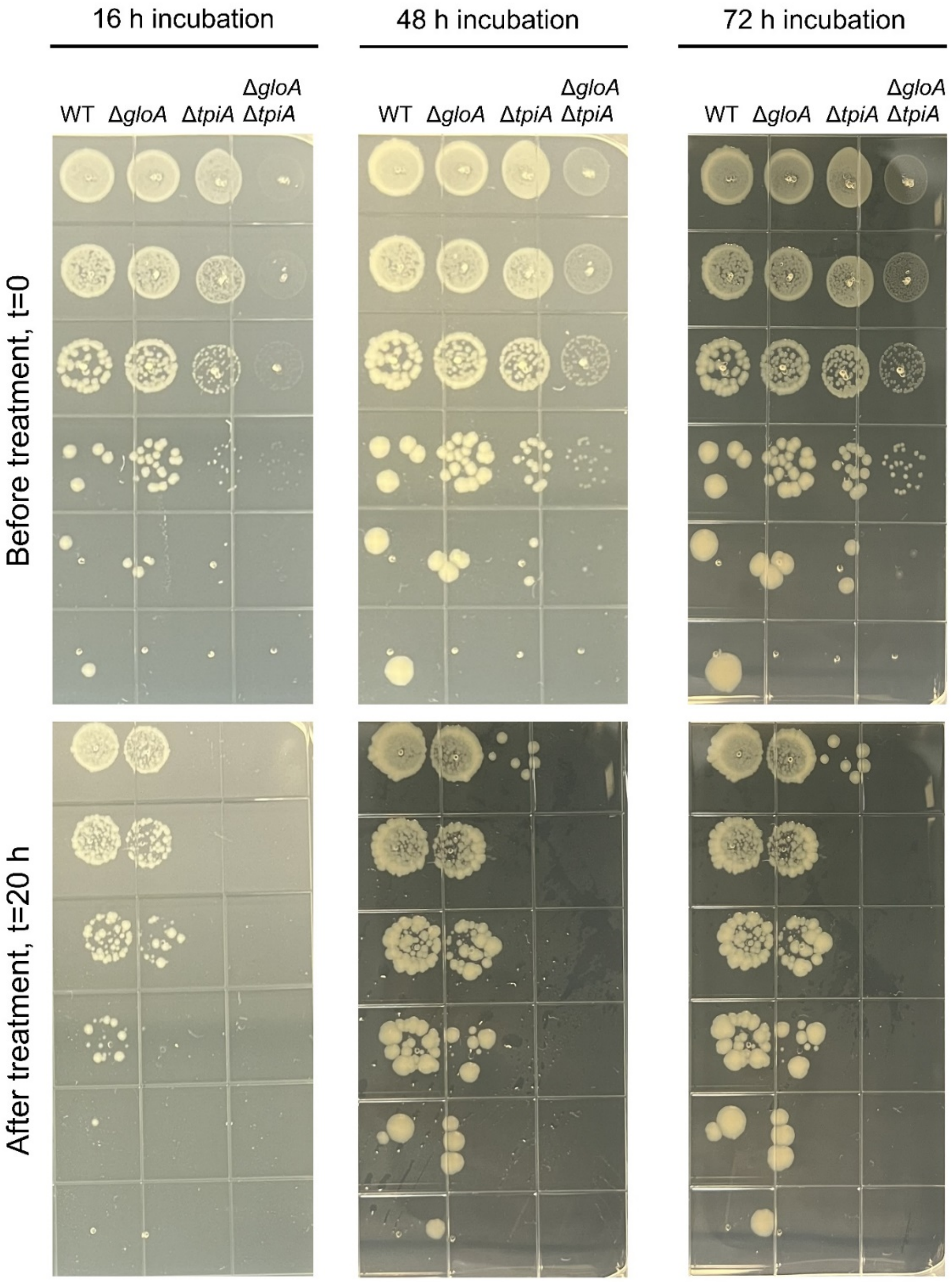


**Fig. S11. Colony formation of WT and mutant strains visualized on agar plates.** Late-stationary-phase (24 h) cultures of *E. coli* WT, Δ*gloA*, Δ*tpiA*, and the double mutant Δ*gloA* Δ*tpiA* were diluted 1:100 into fresh medium and exposed to ofloxacin (5 µg/ml) for 20 h. After the treatment, cells were harvested, washed with 1× PBS, and plated on LB agar to count CFU. Plates were incubated for 72 h to allow small colonies to develop fully, and images were captured at specified time points. N=4.

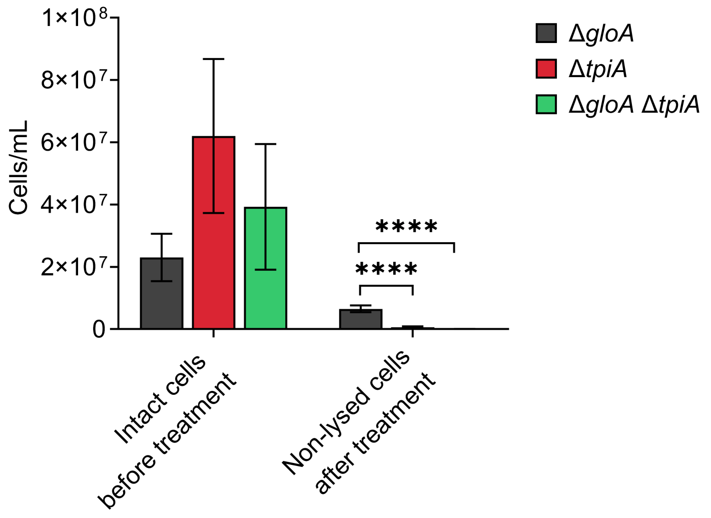


**Fig. S12. Flow cytometry quantification of intact cells after ampicillin treatment.** The fraction of intact (GFP-positive) cells was measured by flow cytometry before and after 20 h of ampicillin treatment in WT, Δ*gloA*, Δ*tpiA*, and Δ*gloA*Δ*tpiA* strains. N = 4. Statistical analysis was performed using two-way ANOVA with Dunnett's multiple comparisons test; ****$P < 0.0001$. Data represent mean ± standard deviation from four independent biological replicates.

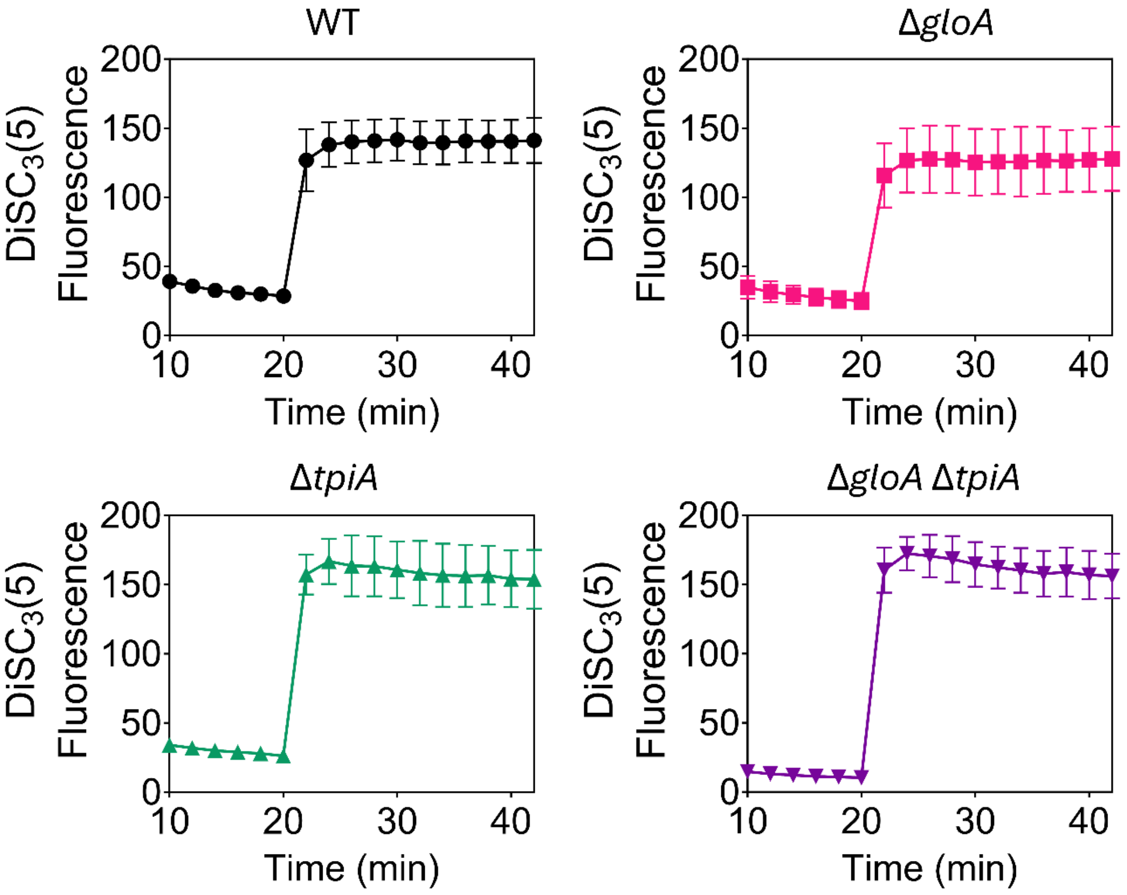


**Fig. S13. Control experiment for the DiSC$_3$(5) assay.** Late stationary phase (24 h) cells of *E. coli* WT, Δ*gloA*, Δ*tpiA*, and the double mutant Δ*gloA* Δ*tpiA* were diluted to an $OD_{600}$ of 0.1, washed with buffer containing 5 mM HEPES and 20 mM glucose, and stained with 1 µM DiSC$_3$(5) dye. After reaching equilibrium, polymyxin B was added, and fluorescence was measured at the indicated time points using a plate reader. Polymyxin B was used as a control to confirm proper dye function. N = 4. Data represent the mean ± standard deviation.

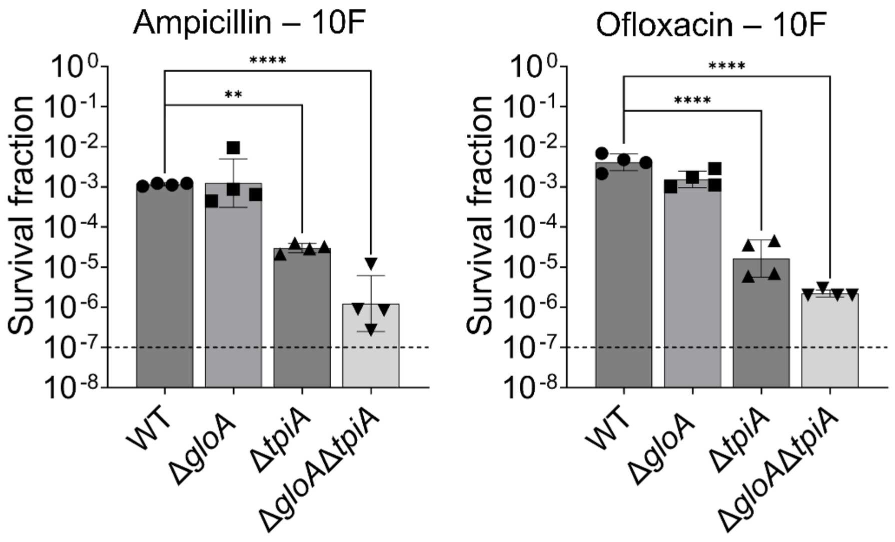


**Fig. S14. Assessment of survival fraction at increased cell density.** *E. coli* WT and mutant strains (Δ*gloA*, Δ*tpiA*, and the double mutant Δ*gloA* Δ*tpiA*) at late stationary phase (24 h) were diluted 1:10-fold and immediately treated with ampicillin (200 µg/ml) and ofloxacin (5 µg/ml) for 20 h. After the treatment, cells were collected, washed with 1× PBS to remove the antibiotics and plated on LB agar to count CFU levels. N=4. Statistical analysis was performed between the WT and mutant strains using one-way ANOVA with Dunnett's post-test. $^{*}P<0.05$, $^{**}P < 0.01$, $^{***}P < 0.001$, and $^{****}P < 0.0001$. Data represent the mean ± standard deviation.

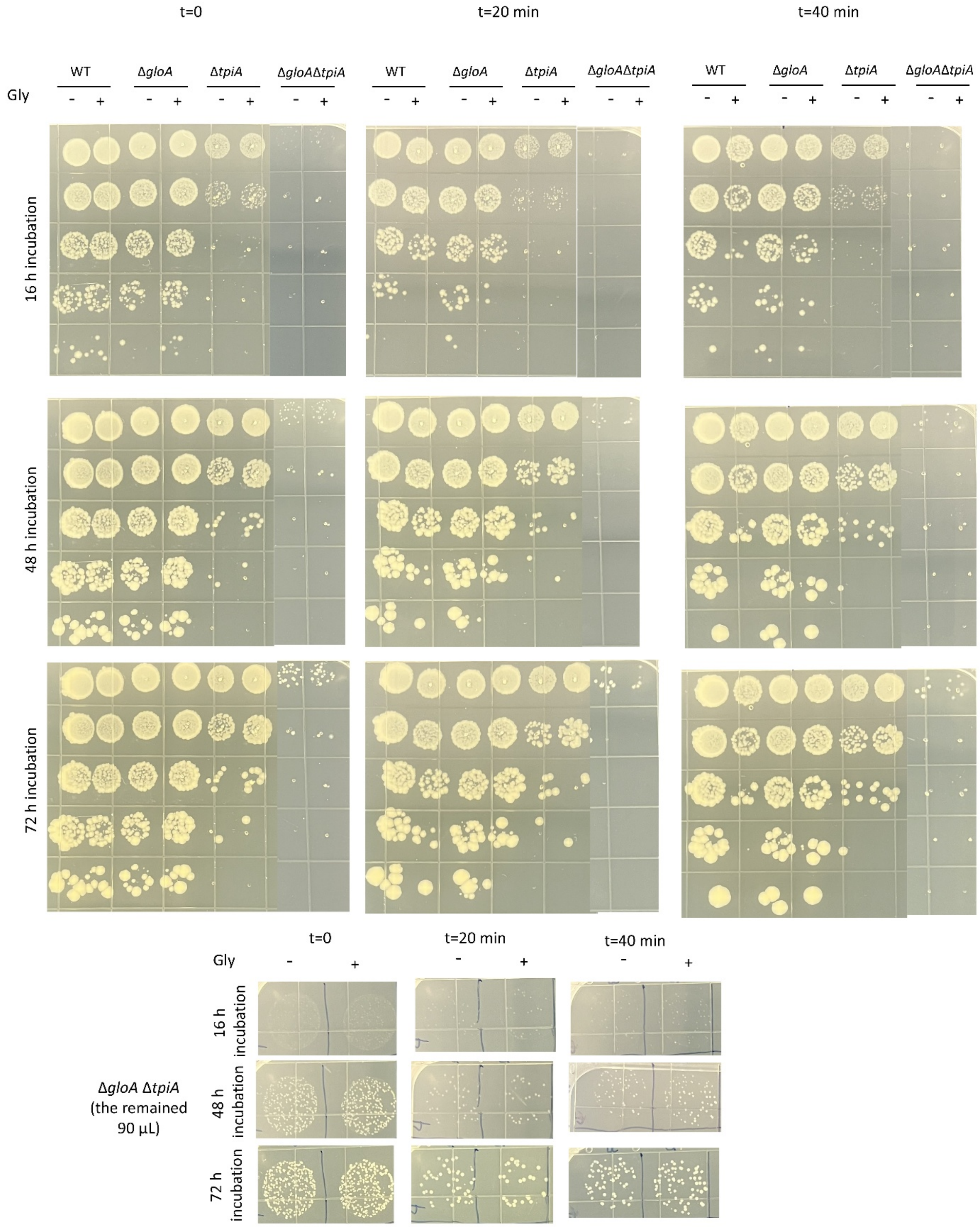


**Fig. S15. Colony formation resulting from the aminoglycoside potentiation assay for WT and mutant strains, visualized on agar plates.** *E. coli* WT and mutant strains (Δ*gloA*, Δ*tpiA*, and the double mutant Δ*gloA* Δ*tpiA*) at late stationary phase (24 h) were diluted 1:10 and immediately exposed to **ampicillin** (200 µg/ml) for 20 hours. After treatment, surviving cells were collected, washed to remove residual antibiotics, and then treated with either DI water or 60 mM glycerol. Kanamycin (25 µg/ml) was subsequently added to designated cultures. At specified time points, 100 µL of cell suspension was collected, washed with 1× PBS, and resuspended in 100 µL of PBS. A 10 µL aliquot of each suspension was plated on LB agar to determine CFU levels. The remaining 90 µL of the double mutant suspension was also plated to increase the limit of CFU detection. Plates were incubated for 72 hours to allow full development of small colonies, and images were taken at the indicated time points. N=4.

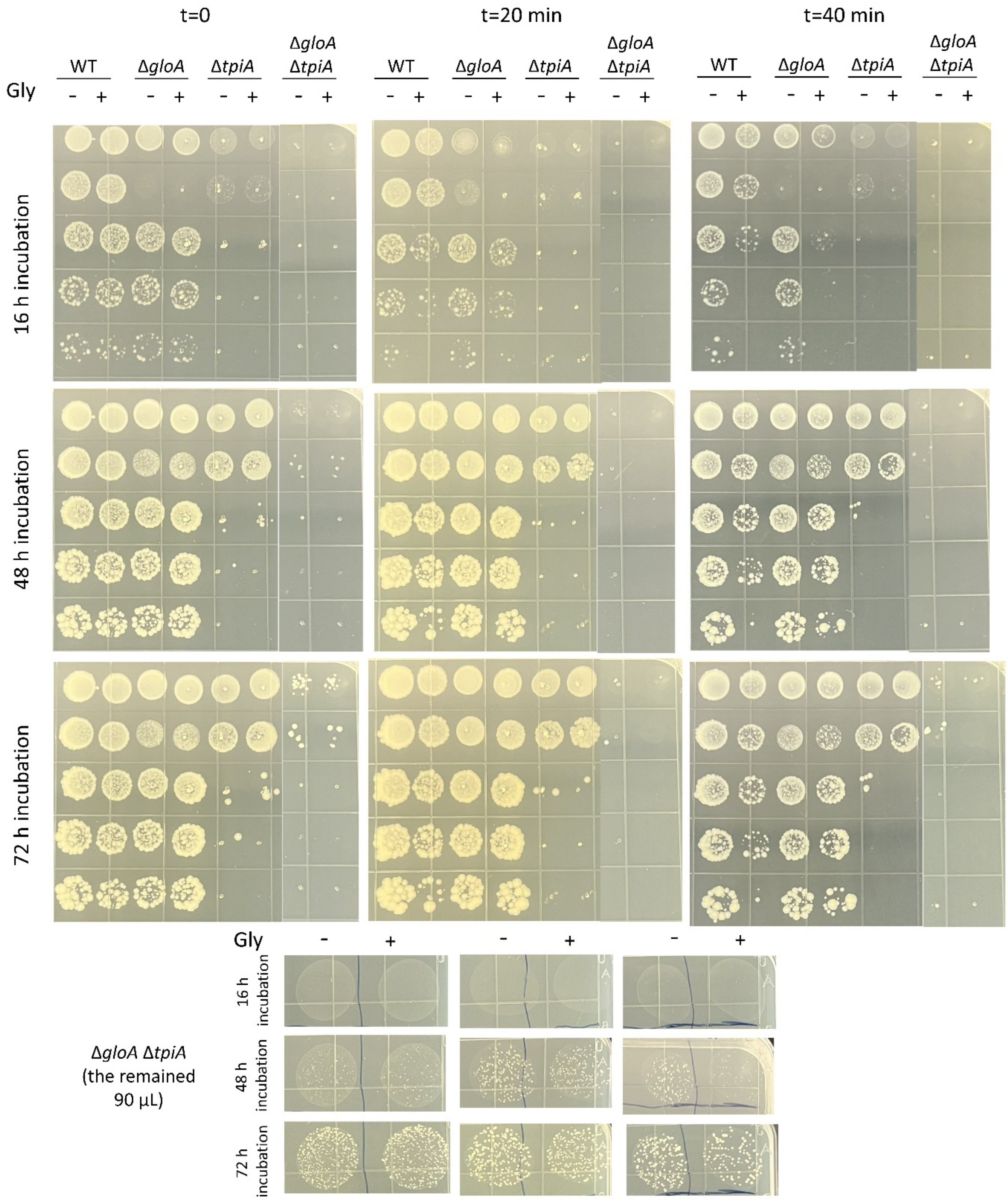


**Fig. S16. Colony formation resulting from the aminoglycoside potentiation assay for WT and mutant strains, visualized on agar plates.** *E. coli* WT and mutant strains (Δ*gloA*, Δ*tpiA*, and the double mutant Δ*gloA* Δ*tpiA*) at late stationary phase (24 h) were diluted 1:10 and immediately exposed to **ofloxacin** (5 µg/ml) for 20 hours. After treatment, surviving cells were collected, washed to remove residual antibiotics, and then treated with either DI water or 60 mM glycerol.

Kanamycin (25 μg/ml) was subsequently added to designated cultures. At specified time points, 100 μL of cell suspension was collected, washed with 1× PBS, and resuspended in 100 μL of PBS. A 10 μL aliquot of each suspension was plated on LB agar to determine CFU levels. The remaining 90 μL of the double mutant suspension was also plated to increase the limit of CFU detection. Plates were incubated for 72 hours to allow full development of small colonies, and images were taken at the indicated time points. N=4.

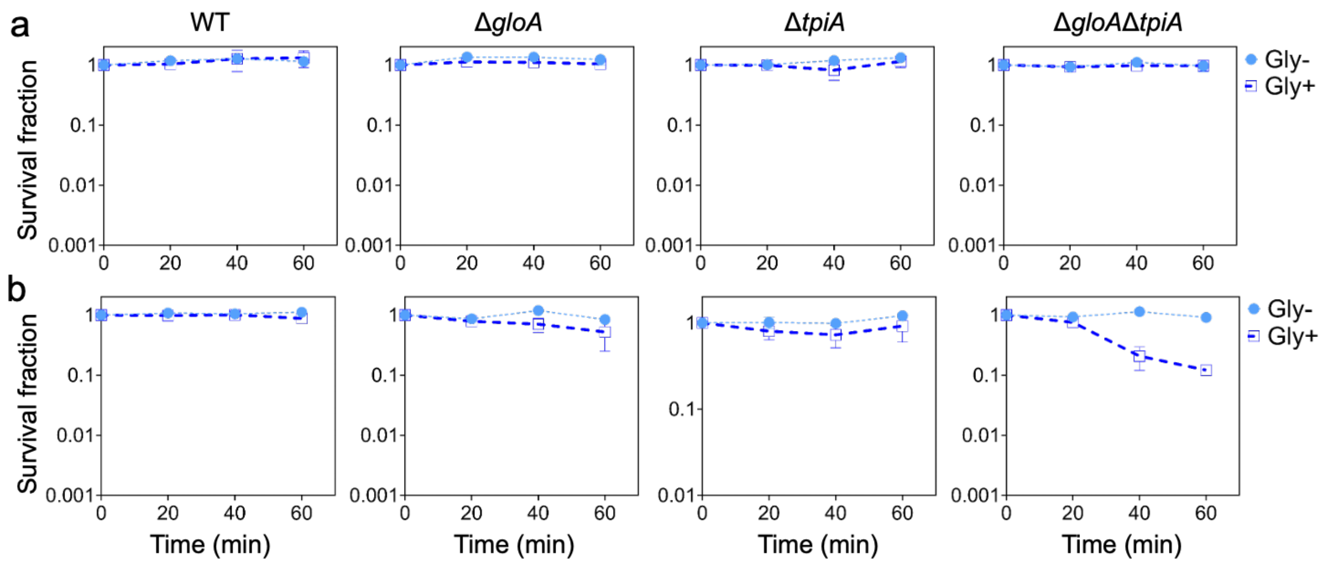


**Fig. S17. Recovery assays of antibiotic-tolerant persister cells in the absence of aminoglycosides.** Late stationary-phase cultures were treated with ampicillin (panel a) or ofloxacin (panel b) for 20 h, then incubated in M9 minimal medium supplemented with either glycerol (Gly+) (60 mM) or deionized water (glycerol negative, Gly−) as a control. N = 4.

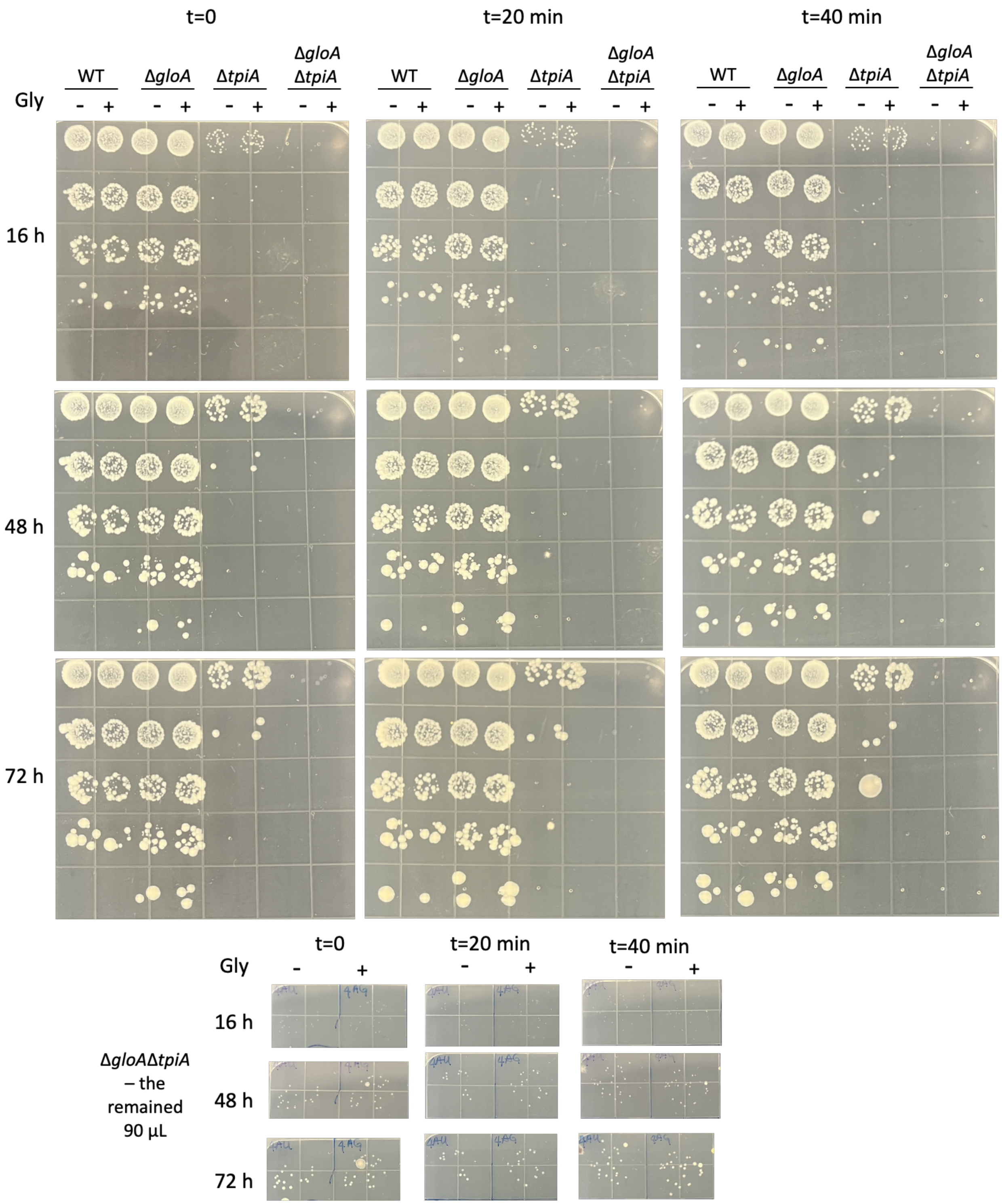


**Fig. S18. Colony formation from the recovery assays, visualized on agar plates.** *E. coli* WT and mutant strains (Δ*gloA*, Δ*tpiA*, and the double mutant Δ*gloA* Δ*tpiA*) at late stationary phase (24 h) were diluted 1:10 and immediately exposed to **ampicillin** (200 µg/ml) for 20 hours. After treatment, surviving cells were collected, washed to remove residual antibiotics, and then treated with either DI water or 60 mM glycerol. **No kanamycin was added**. At specified time points, 100 μL of cell suspension was collected, washed with 1× PBS, and resuspended in 100 μL of PBS. A 10 μL aliquot of each suspension was plated on LB agar to determine CFU levels. The remaining 90 μL of the double mutant suspension was also plated to increase the limit of CFU detection. Plates were incubated for 72 hours to allow full development of small colonies, and images were taken at the indicated time points. N=4.

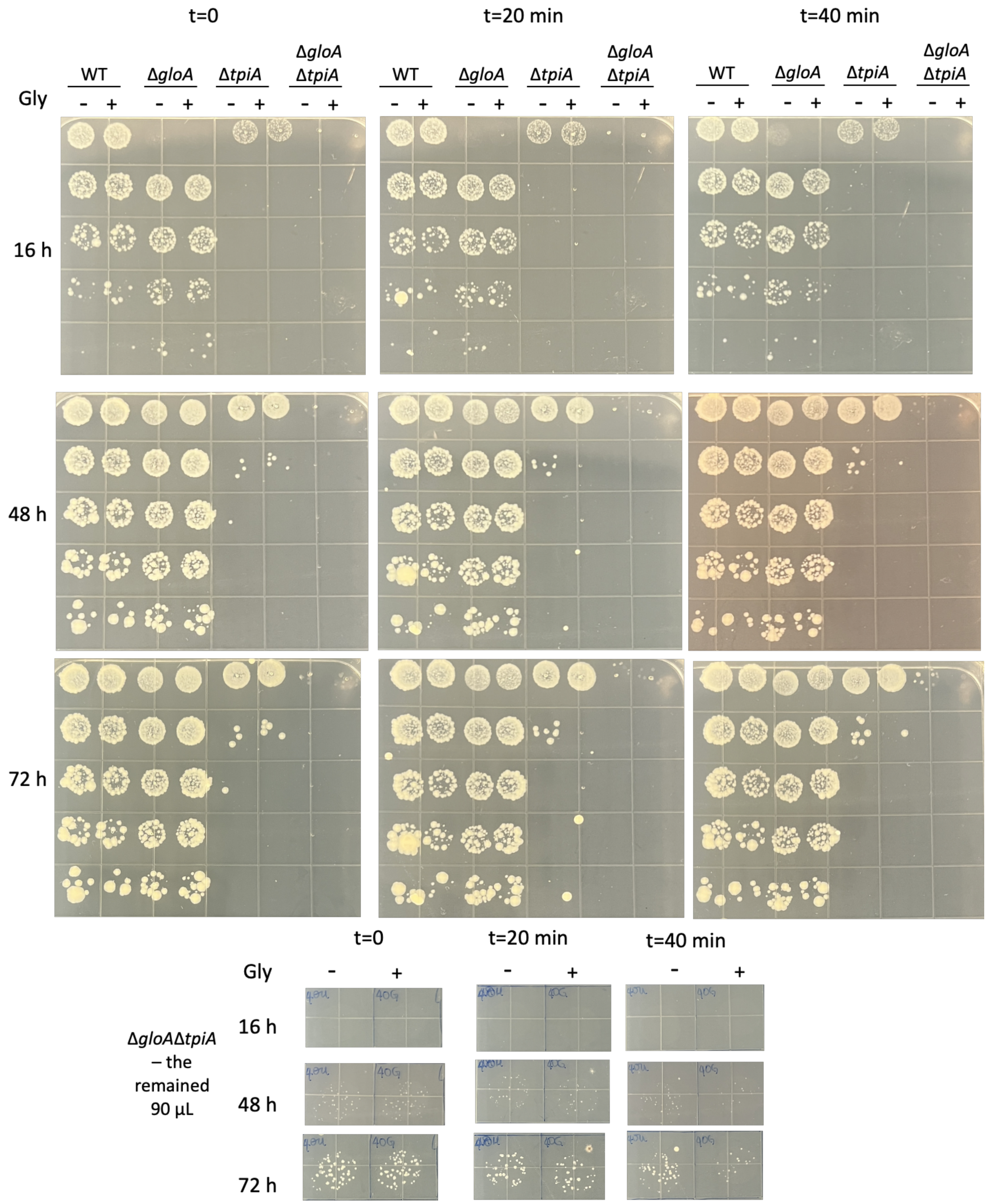


**Fig. S19. Colony formation from the recovery assays, visualized on agar plates.** *E. coli* WT and mutant strains (Δ*gloA*, Δ*tpiA*, and the double mutant Δ*gloA* Δ*tpiA*) at late stationary phase (24 h) were diluted 1:10 and immediately exposed to **ofloxacin** (5 µg/ml) for 20 hours. After treatment, surviving cells were collected, washed to remove residual antibiotics, and then treated with either DI water or 60 mM glycerol. **No kanamycin was added**. At specified time points, 100 μL of cell suspension was collected, washed with 1× PBS, and resuspended in 100 μL of PBS. A 10 μL aliquot of each suspension was plated on LB agar to determine CFU levels. The remaining 90 μL of the double mutant suspension was also plated to increase the limit of CFU detection. Plates were incubated for 72 hours to allow full development of small colonies, and images were taken at the indicated time points. N=4.

### Supplementary Tables

**Table S1. Functions of proteins are found to be upregulated or downregulated based on proteomics analysis.**

| Gene | Upregulated/ Downregulated | Functions | Note |
|---|---|---|---|
| *comR* | Upregulated | DNA-binding transcriptional repressor inhibited by the CRP/cAMP complex and involved in reactions with $Cu^{2+}$ | (Duarte-Velázquez et al., 2022; Mermod et al., 2012) |
| *yhdE* | Upregulated | Nucleoside triphosphate pyrophosphatase that preferentially hydrolyzes UTP and dTTP. Its role is crucial for maintaining phosphate balance and driving various biochemical reactions, including RNA and DNA synthesis | (Jin et al., 2015; Tchigvintsev et al., 2013) |
| *ompX* | Upregulated | Outer membrane protein involved in secreting extracellular proteins. Studies found that an *ompX* mutant exhibited impaired motility and increased EPS production | (Otto et al., 2001; Otto & Hermansson, 2004) |
| *ydcL* | Upregulated | DUF3313 domain-containing lipoprotein that attaches to the cell membrane and bridges proteins and the membrane, playing important roles in nutrient uptake, adhesion, and signal transduction | (Juncker et al., 2003) |
| *arcB* | Upregulated | Sensor histidine kinase that mediates anaerobic repression of numerous enzymes associated with aerobic metabolism | (Iuchi et al., 1989; Iuchi & Lin, 1992) |
| *ndk* | Upregulated | Nucleoside diphosphate kinase that facilitates phosphate transfer from NTPs to NDPs to produce different nucleoside triphosphates, crucial for | (Roisin & Kepes, 1978) |

| | | | |
|---|---|---|---|
| | | maintaining nucleotide pools during nucleic acid synthesis | |
| *fic* | Upregulated | Stationary-phase protein and putative adenosine monophosphate-protein transferase that acts as a cellular energy sensor and regulator, activated when energy levels are low and switching the cell to energy-producing mode | (Utsumi et al., 1982) |
| *rclA* | Upregulated | Cupric reductase that mediates oxidative stress through NADH-dependent cupric reductase and hypothiocyanite reductase activities | (Baek et al., 2020; Derke et al., 2020; Meredith et al., 2022) |
| *fepB* | Upregulated | Ferric enterobactin ABC transporter periplasmic binding protein that transports $Fe^{2+}$ into the cell | (Sprencel et al., 2000) |
| *btuE* | Upregulated | Thioredoxin/glutathione peroxidase that protects cells from oxidative stress | (Arenas et al., 2010) |
| *rplD* | Upregulated | 50S ribosomal subunit protein L4 | (Freedman et al., 1987) |
| *ydhS* | Upregulated | FAD/NAD(P) binding domain-containing protein; NADP is used in anabolic processes for lipid and nucleic acid synthesis | (Keseler et al., 2011) |
| *yqcC* | Upregulated | DUF446 domain-containing protein involved in biofilm formation | (Keseler et al., 2011) |
| *dkgB* | Upregulated | methylglyoxal reductase that reduces the toxic compound methylglyoxal into lactaldehyde | (Di Luccio et al., 2006) |
| *yahK* | Upregulated | NADPH-dependent aldehyde reductase involved in biofilm formation in reduced-genome *E. coli* | (Pick et al., 2013) |
| *iaaA* | Upregulated | Isoaspartyl dipeptidase proenzyme | (Keseler et al., 2011) |

| | | | |
|---|---|---|---|
| *mfd* | Downregulated | Transcription-repair coupling factor that facilitates the repair of lesions on the template strand | (Selby et al., 1991) |
| *alsB* | Downregulated | D-allose ABC transporter periplasmic binding protein related to the glycolysis pathway because *E. coli* produces D-allose from D-glucose | (Keseler et al., 2011) |
| *ybgI* | Downregulated | Radiation resistance protein | (Sergeeva et al., 2018) |
| *ulaD* | Downregulated | 3-keto-L-gluconate-6-phosphate decarboxylase induced by CRP/cAMP and involved in anaerobic L-ascorbate degradation | (Yew & Gerlt, 2002) |
| *asnS* | Downregulated | Asparagine-tRNA ligase | (Yamamoto et al., 1977) |
| *frdB* | Downregulated | Fumarate reductase [Fe-S] protein and a crucial enzyme in anaerobic respiration; it catalyzes the reduction of fumarate to succinate, a key step in energy generation when oxygen and other electron acceptors are unavailable | (Cole et al., 1982) |
| *pepT* | Downregulated | Tripeptidase that cleaves amino-terminal leucine, lysine, methionine, and phenylalanine residues from certain tripeptides; its expression is upregulated during biofilm development and anaerobic growth | (Sussman & Gilvarg, 1970) |
| *aspS* | Downregulated | Aspartate-tRNA ligase | (Eriani et al., 1990) |
| *yagU* | Downregulated | DUF1440 domain-containing inner membrane protein that contributes to acid resistance | (Stenberg et al., 2005) |
| *nagB* | Downregulated | Glucosamine-6-phosphate deaminase that generates products entering glycolysis directly | (Holmes & Russell, 1972) |

| | | | |
|---|---|---|---|
| *fabI* | Downregulated | Enoyl-[acyl-carrier-protein] reductase that catalyzes reactions in fatty acid biosynthesis and is essential for growth | (Bergler et al., 1994) |
| *frdA* | Downregulated | Fumarate reductase flavoprotein subunit | (Cole, 1982) |
| *gabD* | Downregulated | Succinate-semialdehyde dehydrogenase (NADP+) induced by CRP/cAMP; it catalyzes reactions producing intermediates for the TCA cycle | (Cozzani et al., 1980) |
| *pykF* | Downregulated | Pyruvate kinase 1, a key allosteric enzyme in glycolysis produces substrates for glycolysis and the TCA cycle | (Kornberg & Malcovati, 1973) |
| *glpA* | Downregulated | Anaerobic glycerol-3-phosphate dehydrogenase subunit A induced by CRP/cAMP, part of the glycerol degradation I pathway that converts G3P to DHAP using fumarate as the terminal electron acceptor under anaerobic conditions | (Cole et al., 1988; Schryvers & Weiner, 1982) |
| *kdsD* | Downregulated | D-arabinose 5-phosphate isomerase and a constituent of the cell envelope lipopolysaccharide | (Lim & Cohen, 1966) |
| *degP* | Downregulated | Periplasmic serine endoprotease required for survival at high temperature; it degrades abnormal, oxidatively damaged, and aggregated proteins in the periplasm | (Strauch et al., 1989) |
| *aspA* | Downregulated | Aspartate ammonia-lyase induced by CRP/cAMP and expressed under both aerobic and anaerobic conditions | (Rudolph & Fromm, 1971) |
| *ldhA* | Downregulated | D-lactate dehydrogenase is specific for producing D-lactate | (Tarmy & Kaplan, 1968) |

| | | | |
|---|---|---|---|
| *gabT* | Downregulated | 4-aminobutyrate aminotransferase induced by CRP/cAMP and involved in L-lysine degradation | (Dover & Halpern, 1974) |
| *gnd* | Downregulated | 6-phosphogluconate dehydrogenase is involved in the pentose phosphate pathway | (Fraenkel, 1968) |
| *dcd* | Downregulated | dCTP deaminase catalyzes the deamination of dCTP to dUTP, a key step in dTTP biosynthesis and pyrimidine deoxyribonucleotide synthesis | (O'Donovan et al., 1971) |
| *glpB* | Downregulated | Anaerobic glycerol-3-phosphate dehydrogenase subunit B induced by CRP/cAMP, part of the glycerol degradation pathway | (Cole et al., 1988) |
| *nagE* | Downregulated | N-acetylglucosamine-specific PTS enzyme II, part of the sugar transport phosphotransferase system, enters glycolysis | (White, 1970) |
| *hemE* | Downregulated | Uroporphyrinogen decarboxylase is involved in heme B biosynthesis | (Săsărman et al., 1975) |
| *poxB* | Downregulated | Pyruvate oxidase, a peripheral membrane enzyme catalyzing oxidative decarboxylation of pyruvate to acetate and $CO_2$; this reaction is coupled to the ETC via ubiquinone | (Abdel-Hamid et al., 2001) |
| *thiL* | Downregulated | Thiamine monophosphate kinase, is involved in thiamine pyrophosphate (TPP) biosynthesis | (Imamura & Nakayama, 1982) |
| *sufD* | Downregulated | Fe-S cluster scaffold complex subunit required for iron acquisition | (Saini et al., 2010) |
| *pykA* | Downregulated | Pyruvate kinase 2, part of the glycolysis pathway | (Kornberg & Malcovati, 1973) |

| | | | |
|---|---|---|---|
| *uspF* | Downregulated | Universal stress protein that promotes adhesion at the expense of motility | (Saveanu et al., 2002) |
| *ydjA* | Downregulated | Putative oxidoreductase | (Choi et al., 2008) |
| *ptsG* | Downregulated | Glucose-specific PTS enzyme IIBC component induced by CRP/cAMP, involved in glycolysis and mediates glucose uptake and phosphorylation | (Keseler et al., 2011) |
| *yqjI* | Downregulated | DNA-binding transcriptional repressor | (Blahut et al., 2018) |
| *gapA* | Downregulated | Glyceraldehyde-3-phosphate dehydrogenase induced by CRP/cAMP, part of gluconeogenesis and glycolysis | (Hillman, 1979) |
| *ackA* | Downregulated | Acetate kinase | (Hesslinger et al., 1998) |
| *glpT* | Downregulated | sn-glycerol 3-phosphate:phosphate antiporter induced by CRP/cAMP, part of the uptake system for sn-G3P | (Rao et al., 1993) |

**Table S2. Survival fractions of knockout strains following ampicillin and ofloxacin treatment.**

| KO strain | Ampicillin survival fraction | | Ofloxacin survival fraction |
|---|---|---|---|
| *ΔrbfA* | 2.94E-08 | | 1.89E-06 |
| *ΔaspA* | 3.45E-08 | | 1.52E-06 |
| *ΔlysS* | 3.70E-08 | | 2.27E-07 |
| *Δefp* | 4.35E-08 | | 9.33E-07 |
| *ΔselB* | 4.55E-08 | | 1.74E-06 |
| *ΔglpT* | 5.26E-08 | | 5.00E-07 |
| *ΔsdhA* | 1.25E-07 | | 8.57E-07 |
| *ΔglpA* | 1.30E-07 | | 9.13E-07 |
| *ΔcarB* | 1.67E-07 | | 5.50E-06 |

| | | | |
|---|---|---|---|
| *ΔpolA* | 2.00E-07 | | 7.37E-07 |
| *ΔyahK* | 3.87E-07 | | 1.92E-06 |
| *ΔyhdE* | 8.15E-07 | | 4.78E-07 |
| *ΔclpA* | 1.88E-06 | | 3.83E-06 |
| *ΔompW* | 8.64E-06 | | 4.40E-05 |
| *ΔgroL* | 1.14E-05 | | 5.26E-05 |
| *ΔlacI* | 1.29E-05 | | 9.67E-05 |
| *ΔasnB* | 1.36E-05 | | 4.71E-07 |
| *ΔlacY* | 1.61E-05 | | 5.60E-05 |
| *ΔyfcH* | 2.94E-05 | | 6.00E-04 |
| *ΔdegP* | 5.71E-05 | | 1.40E-04 |
| *ΔglpB* | 7.50E-05 | | 5.56E-03 |
| *ΔnuoF* | 8.75E-05 | | 2.11E-02 |
| *ΔackA* | 1.15E-04 | | 1.14E-03 |
| *Δspy* | 1.92E-04 | | 1.31E-03 |
| *Δfic* | 3.11E-04 | | 1.82E-02 |
| *ΔdkgB* | 3.14E-04 | | 5.13E-02 |
| *ΔfepB* | 5.67E-04 | | 6.55E-02 |
| *Δgnd* | 6.36E-04 | | 1.10E-02 |
| *ΔglpF* | 6.84E-04 | | 8.89E-03 |
| *ΔilvN* | 7.33E-04 | | 1.92E-02 |
| *ΔbtuE* | 1.06E-03 | | 2.50E-02 |
| *ΔpykA* | 1.13E-03 | | 7.22E-03 |
| *ΔldhA* | 1.13E-03 | | 1.38E-02 |
| *ΔptsG* | 1.42E-03 | | 1.81E-02 |
| *ΔiaaA* | 1.44E-03 | | 2.20E-02 |
| *ΔompX* | 1.52E-03 | | 9.00E-02 |
| *ΔpykF* | 1.67E-03 | | 4.38E-02 |
| *ΔcomR* | 1.67E-03 | | 2.10E-01 |
| *ΔgabD* | 1.83E-03 | | 7.86E-03 |
| *ΔfrdB* | 1.93E-03 | | 1.33E-02 |
| *ΔyqcC* | 3.13E-03 | | 8.46E-02 |
| *Δmfd* | 3.50E-03 | | 1.46E-02 |
| *ΔfrdA* | 3.53E-03 | | 1.06E-02 |
| *Δndk* | 3.64E-03 | | 4.07E-02 |
| *ΔydhS* | 3.79E-03 | | 1.10E-01 |
| *ΔpoxB* | 4.17E-03 | | 3.29E-02 |
| *ΔnuoE* | 4.50E-03 | | 9.75E-03 |
| *ΔydcL* | 4.93E-03 | | 8.67E-02 |

| | | | |
|---|---|---|---|
| *ΔarcB* | 5.22E-03 | | 1.67E-01 |
| *ΔrclA* | 2.43E-02 | | 5.40E-02 |

**Table S3. Differential abundance of major *E. coli* phospholipid species and glycerol-3-phosphate in early and late stationary phase.**

Wild-type *E. coli* cells were grown under stationary-phase culture conditions and harvested at early (t=5 h) and late (t=24 h) stationary phase. Metabolite and lipid abundances were quantified by LC–MS at Metabolon, Inc., and statistical significance was assessed using an ANOVA contrast for the early (ESP) versus late (LSP) stationary phase comparison; q-values reflect Benjamini–Hochberg correction for multiple testing. Note that phosphatidylethanolamine (PE) and phosphatidylglycerol (PG) species are the dominant *E. coli* membrane phospholipids and represent the most abundant potential sources of glycerol-3-phosphate generated through phospholipid turnover. Fold changes represent the LSP/ESP ratio, with values less than 1 indicating reduction during stationary-phase progression. Red: Significantly upregulated; green: significantly downregulated. Raw data for this analysis are available in the Reference (Ngo et al., 2025).

| Sub-pathway | Biochemical name | LSP/ESP | p-value | q-value |
|---|---|---|---|---|
| Phosphatidylethanolamine (PE) | 1,2-dipalmitoyl-GPE (16:0/16:0) | **1.58** | 0.00010000 | 0.00010000 |
| | 1-palmitoyl-2-stearoyl-GPE (16:0/18:0) | **0.75** | 0.02300000 | 0.01010000 |
| | 1-palmitoyl-2-oleoyl-GPE (16:0/18:1) | **0.07** | 0.00000002 | 0.00000005 |
| | 1-palmitoleoyl-2-oleoyl-GPE (16:1/18:1) | **0.02** | 0.00000000 | 0.00000001 |
| | 1-stearoyl-2-oleoyl-GPE (18:0/18:1) | **0.21** | 0.00000039 | 0.00000072 |
| | 1-stearoyl-2-linoleoyl-GPE (18:0/18:2) | **3.03** | 0.00005060 | 0.00004462 |
| | 1,2-dioleoyl-GPE (18:1/18:1) | **0.03** | 0.00000001 | 0.00000003 |
| Phosphatidylglycerol (PG) | 1,2-dipalmitoyl-GPG (16:0/16:0) | **1.76** | 0.00020000 | 0.00020000 |
| | 1-palmitoyl-2-palmitoleoyl-GPG (16:0/16:1) | **0.25** | 0.00000139 | 0.00000200 |
| | 1-palmitoyl-2-oleoyl-GPG (16:0/18:1) | **0.53** | 0.00020000 | 0.00010000 |
| | 1-palmitoleoyl-2-oleoyl-GPG (16:1/18:1) | **0.07** | 0.00000003 | 0.00000008 |
| | 1,2-dioleoyl-GPG (18:1/18:1) | **0.12** | 0.00000019 | 0.00000039 |
| Lysophospholipid | 1-palmitoyl-GPA (16:0) | 1.09 | 0.99650000 | 0.27890000 |
| | 1-palmitoleoyl-GPA (16:1) | **0.28** | 0.00080000 | 0.00050000 |
| | 1-palmitoyl-GPE (16:0) | **0.23** | 0.00000324 | 0.00000415 |
| | 1-stearoyl-GPE (18:0) | **0.44** | 0.00040000 | 0.00030000 |
| | 2-stearoyl-GPE (18:0) | 0.49 | 0.15150000 | 0.05360000 |
| | 1-oleoyl-GPE (18:1) | **0.03** | 0.00000001 | 0.00000003 |
| | 1-palmitoyl-GPG (16:0) | **0.14** | 0.00009467 | 0.00007891 |

| | | | | |
|---|---|---|---|---|
| | 1-stearoyl-GPG (18:0) | 1.33 | 0.49060000 | 0.15460000 |
| | 1-oleoyl-GPG (18:1) | **0.03** | 0.00000000 | 0.00000001 |
| Glycerolipid Metabolism | **glycerol 3-phosphate** | **0.33** | 0.00080000 | 0.00050000 |

**Table S4. Differential abundance of major *E. coli* phospholipid species and glycerol-3-phosphate in phenothiazine-treated stationary-phase cells.**

Wild-type *E. coli* cells were grown under stationary-phase culture conditions, treated with thioridazine (TDZ) at 5 h of growth, and harvested for metabolomic analysis at 24 h. Metabolite and lipid abundances were quantified by LC–MS at Metabolon, Inc., and statistical significance for TDZ-treated versus untreated samples was assessed using the Welch two-sample t-test; q-values reflect Benjamini–Hochberg correction for multiple testing. PE and PG components are the dominant *E. coli* membrane phospholipids and represent the most abundant potential sources of glycerol-3-phosphate generated through phospholipid turnover. Fold changes represent the ratio of TDZ-treated to untreated cells, with values greater than 1 indicating accumulation under PMF-inhibitory conditions. Red: Significantly upregulated; green: significantly downregulated. Raw data for this analysis are available in the Reference (Mohiuddin et al., 2022).

**Note:** Phenothiazine treatment is relevant because these compounds collapse PMF and suppress TCA and ETC activity, creating a defined metabolic state similar to that of our respiratory-deficient mutants. The lipid data associated with this treatment also provide a practical advantage, as they allow assessment of PMF-dependent effects on phospholipid turnover and glycerol-3-phosphate metabolism without requiring exhaustive testing across all metabolic mutants.

| **Sub-pathway** | **Biochemical name** | **TDZ/ Untreated** | **p-value** | **q-value** |
|---|---|---|---|---|
| Phosphatidylethanolamine (PE) | 1,2-dipalmitoyl-GPE (16:0/16:0) | **0.76** | 0.24360000 | 0.10070000 |
| | 1-palmitoyl-2-stearoyl-GPE (16:0/18:0) | **2.43** | 0.00300000 | 0.00190000 |
| | 1-palmitoyl-2-oleoyl-GPE (16:0/18:1) | **16.62** | 0.00030000 | 0.00020000 |
| | 1-palmitoleoyl-2-oleoyl-GPE (16:1/18:1) | **54.02** | 0.07120000 | 0.03290000 |
| | 1-stearoyl-2-oleoyl-GPE (18:0/18:1) | **7.86** | 0.00040000 | 0.00030000 |
| | 1-stearoyl-2-linoleoyl-GPE (18:0/18:2) | **0.33** | 0.00020000 | 0.00020000 |
| | 1,2-dioleoyl-GPE (18:1/18:1) | **40.18** | 0.00450000 | 0.00270000 |
| Phosphatidylglycerol (PG) | 1,2-dipalmitoyl-GPG (16:0/16:0) | **0.56** | 0.00070000 | 0.00050000 |
| | 1-palmitoyl-2-palmitoleoyl-GPG (16:0/16:1) | **4.42** | 0.06970000 | 0.03240000 |
| | 1-palmitoyl-2-oleoyl-GPG (16:0/18:1) | **2.38** | 0.00440000 | 0.00270000 |
| | 1-palmitoleoyl-2-oleoyl-GPG (16:1/18:1) | **16.01** | 0.00040000 | 0.00030000 |
| | 1,2-dioleoyl-GPG (18:1/18:1) | **11.52** | 0.75220000 | 0.26860000 |

| | | | | |
|---|---|---|---|---|
| Lysophospholipid | 1-palmitoyl-GPA (16:0) | 2.88 | 0.23210000 | 0.09690000 |
| | 1-palmitoleoyl-GPA (16:1) | **22.09** | 1.00000000 | 0.33270000 |
| | 1-palmitoyl-GPE (16:0) | **9.17** | 0.00780000 | 0.00450000 |
| | 1-stearoyl-GPE (18:0) | **9.80** | 0.00002107 | 0.00002398 |
| | 2-stearoyl-GPE (18:0) | 1.35 | 0.40530000 | 0.15810000 |
| | 1-oleoyl-GPE (18:1) | **62.35** | 0.00290000 | 0.00180000 |
| | 1-palmitoyl-GPG (16:0) | 3.32 | 0.37180000 | 0.14600000 |
| | 1-stearoyl-GPG (18:0) | 1.19 | 0.00050000 | 0.00040000 |
| | 1-oleoyl-GPG (18:1) | **66.90** | 0.00004725 | 0.00004777 |
| Glycerolipid Metabolism | **glycerol 3-phosphate** | **6.38** | 0.00000732 | 0.00000944 |

**Table S5. Concentrations of bactericidal antibiotics used in persister assays.**

| | Concentration (µg/mL) | |
|---|---|---|
| **Bacterial Strains** | **Ampicillin** | **Ofloxacin** |
| MIC of *E. coli* K-12 MG1655 Wild Type | 1.5-2 | 0.032-0.047 |
| MIC of *E. coli* K-12 MG1655 Δ*gloA* | 1.5-2 | 0.032-0.047 |
| MIC of *E. coli* K-12 MG1655 Δ*tpiA* | 1.5-2 | 0.047-0.064 |
| MIC of *E. coli* K-12 MG1655 Δ*gloA* Δ*tpiA* | 3-4 | 0.094-0.125 |
| Persister Assay Concentration | 200 | 5 |

**Table S6. Bacterial strains and plasmids used in this study.**

| **Bacterial Strains** | **Source** |
|---|---|
| *Escherichia coli* K-12 MG1655 Wild Type | Gift from Dr. Mark P. Brynildsen |
| *Escherichia coli* K-12 BW25113 Wild Type | Keio collection, Cat# OEC4988 |
| *Escherichia coli* K-12 MG1655 Δ*gloA* | This study |
| *Escherichia coli* K-12 MG1655 ΔtpiA | This study |
| *Escherichia coli* K-12 MG1655 ΔgloAΔ*tpiA* | This study |
| *Escherichia coli* K-12 BW25113 Δ*rbfA* | Keio collection, Cat# OEC4988 |
| *Escherichia coli* K-12 BW25113 Δ*aspA* | Keio collection, Cat# OEC4988 |
| *Escherichia coli* K-12 BW25113 Δ*lysS* | Keio collection, Cat# OEC4988 |
| *Escherichia coli* K-12 BW25113 Δ*efp* | Keio collection, Cat# OEC4988 |
| *Escherichia coli* K-12 BW25113 Δ*selB* | Keio collection, Cat# OEC4988 |
| *Escherichia coli* K-12 BW25113 Δ*glpT* | Keio collection, Cat# OEC4988 |
| *Escherichia coli* K-12 BW25113 Δ*sdhA* | Keio collection, Cat# OEC4988 |
| *Escherichia coli* K-12 BW25113 Δ*glpA* | Keio collection, Cat# OEC4988 |
| *Escherichia coli* K-12 BW25113 Δ*carB* | Keio collection, Cat# OEC4988 |
| *Escherichia coli* K-12 BW25113 Δ*polA* | Keio collection, Cat# OEC4988 |
| *Escherichia coli* K-12 BW25113 Δ*yahK* | Keio collection, Cat# OEC4988 |
| *Escherichia coli* K-12 BW25113 Δ*yhdE* | Keio collection, Cat# OEC4988 |

| | |
|---|---|
| *Escherichia coli* K-12 BW25113 Δ*clpA* | Keio collection, Cat# OEC4988 |
| *Escherichia coli* K-12 BW25113 Δ*ompW* | Keio collection, Cat# OEC4988 |
| *Escherichia coli* K-12 BW25113 Δ*groL* | Keio collection, Cat# OEC4988 |
| *Escherichia coli* K-12 BW25113 Δ*lacI* | Keio collection, Cat# OEC4988 |
| *Escherichia coli* K-12 BW25113 Δ*asnB* | Keio collection, Cat# OEC4988 |
| *Escherichia coli* K-12 BW25113 Δ*lacY* | Keio collection, Cat# OEC4988 |
| *Escherichia coli* K-12 BW25113 Δ*yfcH* | Keio collection, Cat# OEC4988 |
| *Escherichia coli* K-12 BW25113 Δ*degP* | Keio collection, Cat# OEC4988 |
| *Escherichia coli* K-12 BW25113 Δ*glpB* | Keio collection, Cat# OEC4988 |
| *Escherichia coli* K-12 BW25113 Δ*nuoF* | Keio collection, Cat# OEC4988 |
| *Escherichia coli* K-12 BW25113 Δ*ackA* | Keio collection, Cat# OEC4988 |
| *Escherichia coli* K-12 BW25113 Δ*spy* | Keio collection, Cat# OEC4988 |
| *Escherichia coli* K-12 BW25113 Δ*fic* | Keio collection, Cat# OEC4988 |
| *Escherichia coli* K-12 BW25113 Δ*dkgB* | Keio collection, Cat# OEC4988 |
| *Escherichia coli* K-12 BW25113 Δ*fepB* | Keio collection, Cat# OEC4988 |
| *Escherichia coli* K-12 BW25113 Δ*gnd* | Keio collection, Cat# OEC4988 |
| *Escherichia coli* K-12 BW25113 Δ*glpF* | Keio collection, Cat# OEC4988 |
| *Escherichia coli* K-12 BW25113 Δ*ilvN* | Keio collection, Cat# OEC4988 |
| *Escherichia coli* K-12 BW25113 Δ*btuE* | Keio collection, Cat# OEC4988 |
| *Escherichia coli* K-12 BW25113 Δ*pykA* | Keio collection, Cat# OEC4988 |
| *Escherichia coli* K-12 BW25113 Δ*ldhA* | Keio collection, Cat# OEC4988 |
| *Escherichia coli* K-12 BW25113 Δ*ptsG* | Keio collection, Cat# OEC4988 |
| *Escherichia coli* K-12 BW25113 Δ*iaaA* | Keio collection, Cat# OEC4988 |
| *Escherichia coli* K-12 BW25113 Δ*ompX* | Keio collection, Cat# OEC4988 |
| *Escherichia coli* K-12 BW25113 Δ*pykF* | Keio collection, Cat# OEC4988 |
| *Escherichia coli* K-12 BW25113 Δ*comR* | Keio collection, Cat# OEC4988 |
| *Escherichia coli* K-12 BW25113 Δ*gabD* | Keio collection, Cat# OEC4988 |
| *Escherichia coli* K-12 BW25113 Δ*frdB* | Keio collection, Cat# OEC4988 |
| *Escherichia coli* K-12 BW25113 Δ*yqcC* | Keio collection, Cat# OEC4988 |
| *Escherichia coli* K-12 BW25113 Δ*mfd* | Keio collection, Cat# OEC4988 |
| *Escherichia coli* K-12 BW25113 Δ*frdA* | Keio collection, Cat# OEC4988 |
| *Escherichia coli* K-12 BW25113 Δ*ndk* | Keio collection, Cat# OEC4988 |
| *Escherichia coli* K-12 BW25113 Δ*ydhS* | Keio collection, Cat# OEC4988 |
| *Escherichia coli* K-12 BW25113 Δ*poxB* | Keio collection, Cat# OEC4988 |
| *Escherichia coli* K-12 BW25113 Δ*nuoE* | Keio collection, Cat# OEC4988 |
| *Escherichia coli* K-12 BW25113 Δ*ydcL* | Keio collection, Cat# OEC4988 |
| *Escherichia coli* K-12 BW25113 Δ*arcB* | Keio collection, Cat# OEC4988 |
| *Escherichia coli* K-12 BW25113 Δ*rclA* | Keio collection, Cat# OEC4988 |
| **Bacterial Plasmids** | **Source or Reference** |
| pUA66-*ftsZ-gfp* | Previous study (Mohiuddin 2022) |

**Table S7. Oligonucleotides for the generation and verification of mutant strains.**

| **Oligonucleotides to Generate Gene Deletions** | | | | | |
|---|---|---|---|---|---|
| **Mutation** | **Forward Primer (5' to 3')** | **Reverse Primer (5' to 3')** | **Source** | | |
| Δ*gloA::Kan* | CGCTATACTAAAACAACATTTTGAATCTGTTAGCCATTTTGAGGATAAAAAGGTGTAGGCTGGAGCTGCTTC | GTAAAGATGCGGGCGCGATGAGTTCACGCCCGGCAGGAGATTAACGGCTGACATGGGAAT | Integrated DNA Technologies, Inc. | | |
| Δ*tpiA::Kan* | GCCATCTTCCTTTATTCGCTTATAAGCGTGGAGAATTAAAGTGTAGGCTGGAGCTGCTTC | GAAAGTAAGTGCCGGATATGAAATCCGGCACCTGTCAGACTTAACGGCTGACATGGGAAT | Integrated DNA Technologies, Inc. | | |
| **Oligonucleotides to Verify Gene Deletions** | | | | | |
| **Mutation** | **External Forward Primer (5' to 3')** | **External Reverse Primer (5' to 3')** | **Internal Forward Primer (5' to 3')** | **Internal Reverse Primer (5' to 3')** | **Source** |
| Δ*gloA::Kan* | TATACCGATTACCCGACGTT | GCTCTTCGTCCAGATCATCC | TTCATACCATGCTGCGCGTT | GACCGGCGTCTTTCTCTTCG | Integrated DNA Technologies, Inc. |
| Δ*tpiA::Kan* | CGCTGTTGAACCGATTAAGC | GCTCTTCGTCCAGATCATCC | AATCGCACCACCGGAAATGT | AAGCCTGTTTAGCCGCTTCTG | Integrated DNA Technologies, Inc. |
